# Gut microbiota dysbiosis is associated with altered tryptophan metabolism and dysregulated inflammatory response in severe COVID-19

**DOI:** 10.1101/2022.12.02.518860

**Authors:** Morgan Essex, Belén Millet Pascual-Leone, Ulrike Löber, Mathias Kuhring, Bowen Zhang, Ulrike Bruening, Raphaela Fritsche-Guenther, Marta Krzanowski, Facundo Fiocca Vernengo, Sophia Brumhard, Ivo Röwekamp, Agata Anna Bielecka, Till Robin Lesker, Emanuel Wyler, Markus Landthaler, Andrej Mantei, Christian Meisel, Sandra Caesar, Charlotte Thibeault, Victor Corman, Lajos Marko, Norbert Suttorp, Till Strowig, Florian Kurth, Leif E. Sander, Yang Li, Jennifer A. Kirwan, Sofia K. Forslund, Bastian Opitz

## Abstract

The clinical course of the 2019 coronavirus disease (COVID-19) is variable and to a substantial degree still unpredictable, especially in persons who have neither been vaccinated nor recovered from previous infection. We hypothesized that disease progression and inflammatory responses were associated with alterations in the microbiome and metabolome. To test this, we integrated metagenome, metabolome, cytokine, and transcriptome profiles of longitudinally collected samples from hospitalized COVID-19 patients at the beginning of the pandemic (before vaccines or variants of concern) and non-infected controls, and leveraged detailed clinical information and post-hoc confounder analysis to identify robust within- and cross-omics associations. Severe COVID-19 was directly associated with a depletion of potentially beneficial intestinal microbes mainly belonging to Clostridiales, whereas oropharyngeal microbiota disturbance appeared to be mainly driven by antibiotic use. COVID-19 severity was also associated with enhanced plasma concentrations of kynurenine, and reduced levels of various other tryptophan metabolites, lysophosphatidylcholines, and secondary bile acids. Decreased abundance of Clostridiales potentially mediated the observed reduction in 5-hydroxytryptophan levels. Moreover, altered plasma levels of various tryptophan metabolites and lower abundances of Clostridiales explained significant increases in the production of IL-6, IFNγ and/or TNFα. Collectively, our study identifies correlated microbiome and metabolome alterations as a potential contributor to inflammatory dysregulation in severe COVID-19.

**Graphical Abstract:** 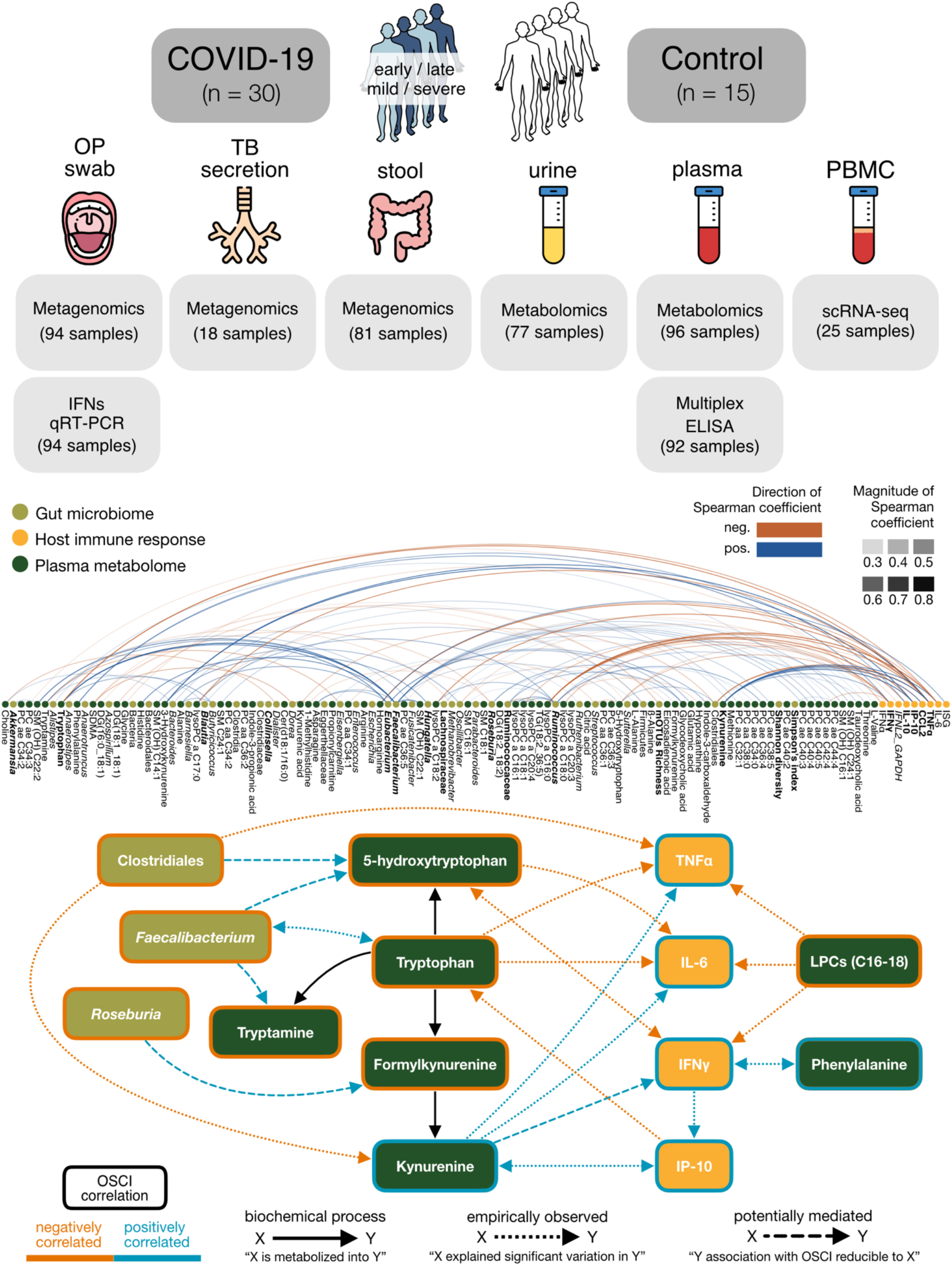

## Introduction

The Coronavirus disease 2019 (COVID-19) pandemic, caused by the severe acute respiratory syndrome coronavirus 2 (SARS-CoV-2), has affected over 600 million individuals and resulted in more than 6 million deaths worldwide by early November 2022 (https://coronavirus.jhu.edu). The infection typically starts with mild to moderate respiratory symptoms. After approximately one week, a minority of infected individuals develop pneumonia which may be complicated by acute respiratory distress syndrome (ARDS), coagulopathy, and multiorgan failure^1,2^. The common kinetics of disease progression together with recent observational studies suggest that disease severity is primarily driven by a dysregulated immune response. Several studies found high levels of proinflammatory cytokines, such as IL-6, TNFα, and IFNγ, as well as T cell lymphopenia, decrease of non-classical (CD14^lo^CD16^hi^) monocytes, and occurrence of neutrophil precursors in the peripheral blood of severe COVID-19 patients^3–7^. Older age, male sex, chronic lung and cardiovascular diseases, diabetes mellitus, obesity, host genetics, and IFN autoantibodies have also been associated with severe disease and death^8–11^, but these factors alone do not appear to explain the wide variability in the clinical course of COVID-19.

Mucosal surfaces of the upper respiratory tract and gut are physiologically colonized with a microbiota that consists of trillions of microbial cells and whose diversity and composition vary widely among individuals^12^. The microbiota constantly generates thousands of unique metabolites that can influence many aspects of human biology^13^. Animal studies have revealed that the microbiota calibrates immune responses during pulmonary and systemic infections, e.g. through production of short-chain fatty acids (SCFAs), tryptophan catabolites, and secondary bile acids^14–17^. Interindividual gut microbiota differences in humans have been associated with variation in cytokine production capacities of peripheral blood cells^18^, and lung microbiota composition has been linked to e.g. baseline levels of proinflammatory cytokines^19^. Previous studies have also indicated an association between COVID-19 status and/or severity, gut microbiota composition, and production of some inflammatory mediators^20–22^. Moreover, a reduced abundance of upper respiratory tract commensals in severe COVID-19 patients has been described^23–25^.

In this study, we longitudinally collected samples from COVID-19 patients with varying disease severity as well as from uninfected controls, and applied an integrated systems approach to characterize the interplay between the microbiome, metabolome and immune system. Using linear mixed-effect models and exhaustive confounder testing to account for clinical and host factors wherever possible, we identify distinct microbiome, metabolome, and immunological signatures of severe COVID-19, and reveal associations between specific members of the microbiota, circulating metabolites, and systemic inflammatory mediators.

## Results

The present work includes a subset of patients enrolled between March and June 2020 in the Pa-COVID-19 cohort, a prospective observational cohort study of patients with COVID-19 at Charité Universitätsmedizin Berlin^26^. We longitudinally collected plasma, stool, urine, and oropharyngeal (OP) swabs from a total of 30 laboratory-confirmed, hospitalized COVID-19 patients with varying degrees of disease severity, as well as 15 uninfected, age- and sex-matched controls (Fig. 1). In parallel, we obtained comprehensive clinical information including underlying diseases, medication before and during hospitalization, and the development of secondary infections (Table 1 and Table S1). We classified patient samples into early or late observation groups based on the number of days since symptom onset (≤10 days or >10 days, respectively). According to the WHO ordinal scale of clinical improvement (OSCI, www.who.int/publications/i/item/covid-19-therapeutic-trial-synopsis), 22 patients (73.3%) had mild disease (maximum OSCI score 1-4), and 8 (26.7%) had a more severe disease course (maximum OSCI score 5-8), 3 of whom died in the hospital. The median duration of hospitalization was 8.5 days (range 3-132 days) excluding patients who died (3). Peripheral blood mononuclear cells (PBMCs) were obtained from 14 patients at an early phase of infection as well as 11 controls, and tracheobronchial secretions (TBS) were collected from 4 ventilated COVID-19 patients. We performed whole metagenome sequencing of stool, OP and TBS samples, metabolomics of plasma and urine, single cell RNA sequencing (scRNAseq) of PBMCs, multiplex cytokine ELISA of plasma, and *IFN* qRT-PCRs of OP samples. Our integrated statistical approach enabled us to analyze -omics and clinical data individually and in conjunction with one another while accounting for a range of potential confounders.

**Table 1:** Clinical characteristics.

| Characteristics | Patients | Controls |
| --- | --- | --- |
| N | 30 | 15 |
| Average age, y (SD) | 58.17 (18.15) | 52.80 (19.37) |
| Female, % (n) | 43.3 (13) | 40.0 (6) |
| Average BMI, (SD) | 26.85 (4.91) | 23.94 (3.52) |
| Active smokers, % (n) | 10.0 (3) | 6.7 (1) |
| Comorbidities |  |  |
| Diabetes mellitus, % (n) | 16.7 (5) | 6.7 (1) |
| Cardiovascular disease, % (n) | 53.3 (16) | 26.7 (4) |
| Chronic lung disease, % (n) | 30.0 (9) | 0 (0) |
| Chronic kidney disease, % (n) | 33.3 (10) | 0 (0) |
| Chronic liver disease, % (n) | 16.7 (5) | 6.7 (1) |
| IBD, % (n) | 3.3 (1) | 0 (0) |
| Dyslipidemia, % (n) | 16.7 (5) | 13.3 (2) |
| Active neoplasia, % (n) | 3.3 (1) | 0 (0) |
| Altered thyroid hormones, % (n) | 23.3 (7) | 6.7 (1) |
| Charlson Comorbidity Index, median (IQR) | 2.5 (4.0) | 1.0 (3.25) |
| OSCI, median (IQR) | 4.0 (2.0) | 0 (0) |
| Gastrointestinal symptoms at admission, % (n) | 23.3 (7) | 0 (0) |
| Medication, median (IQR) | 2 (3.0) | 0 (2.0) |

**Figure 1:**
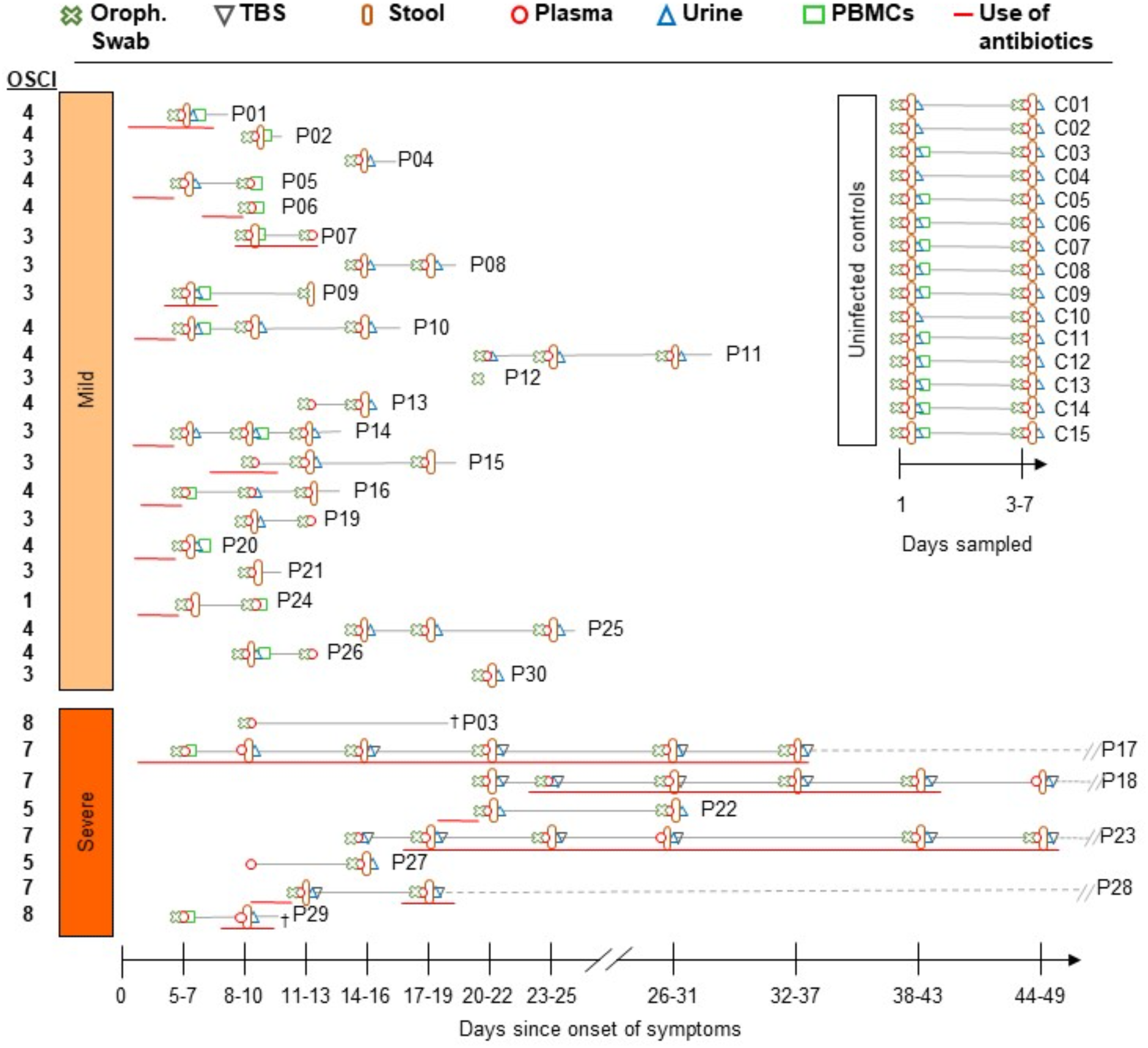
Cohort description and sampling timepoints. Uninfected controls and enrolled patients classified by maximum OSCI score. We refer to scores between 1-4 as mild and scores between 5-8 as severe disease in further discussion. Sampling time points are represented according to the days after symptom onset for patients. The observation and hospitalization period is marked with a solid line, or a dashed gray line when prolonged. Sample materials included oropharyngeal swabs, plasma, peripheral mononuclear blood cells (PBMCs), urine, stool and tracheobronchial secretions (TBS). The use of antibiotics shortly before or during the sampling period is marked for each participant. All control subjects were antibiotic-free for at least 3 months before and during the sampling period.

### Airway and intestinal microbiota disturbance in severe COVID-19

To characterize the microbiota of our cohorts, shotgun metagenomic sequencing was conducted on a total of 94 OP swabs, 18 TBS, and 81 stool samples. The gut microbiota of COVID-19 patients exhibited significantly decreased taxonomic diversity and richness when compared to uninfected controls (FDR-adjusted P<0.001) regardless of infection timepoint, which is in line with previous observations^20,27,28^. These depletions were most strongly associated with disease severity as measured by OSCI score, and negatively correlated with the number of days patients were hospitalized (Fig. 2A, C, Fig. S1). Taxonomic profiling further revealed that severe courses of COVID-19 robustly associated with lower abundances of several commensals including Ruminococcaceae, Lachnospiraceae, *Bifidobacterium, Faecalibacterium, Roseburia*, and *Intestinibacter* in the gut (Fig. 2E, Fig. S1). Importantly, disease severity explained more significant variation in the intestinal abundances of these taxa than antibiotic intake during the sampling period. The depletion of commensals was exacerbated and in some cases potentially mediated by longer hospitalization times. Patients with longer hospitalization times additionally displayed depletions of e.g. *Rothia* and *Actinomyces* along with increased colonization by *Pseudomonas*.

**Figure 2:**
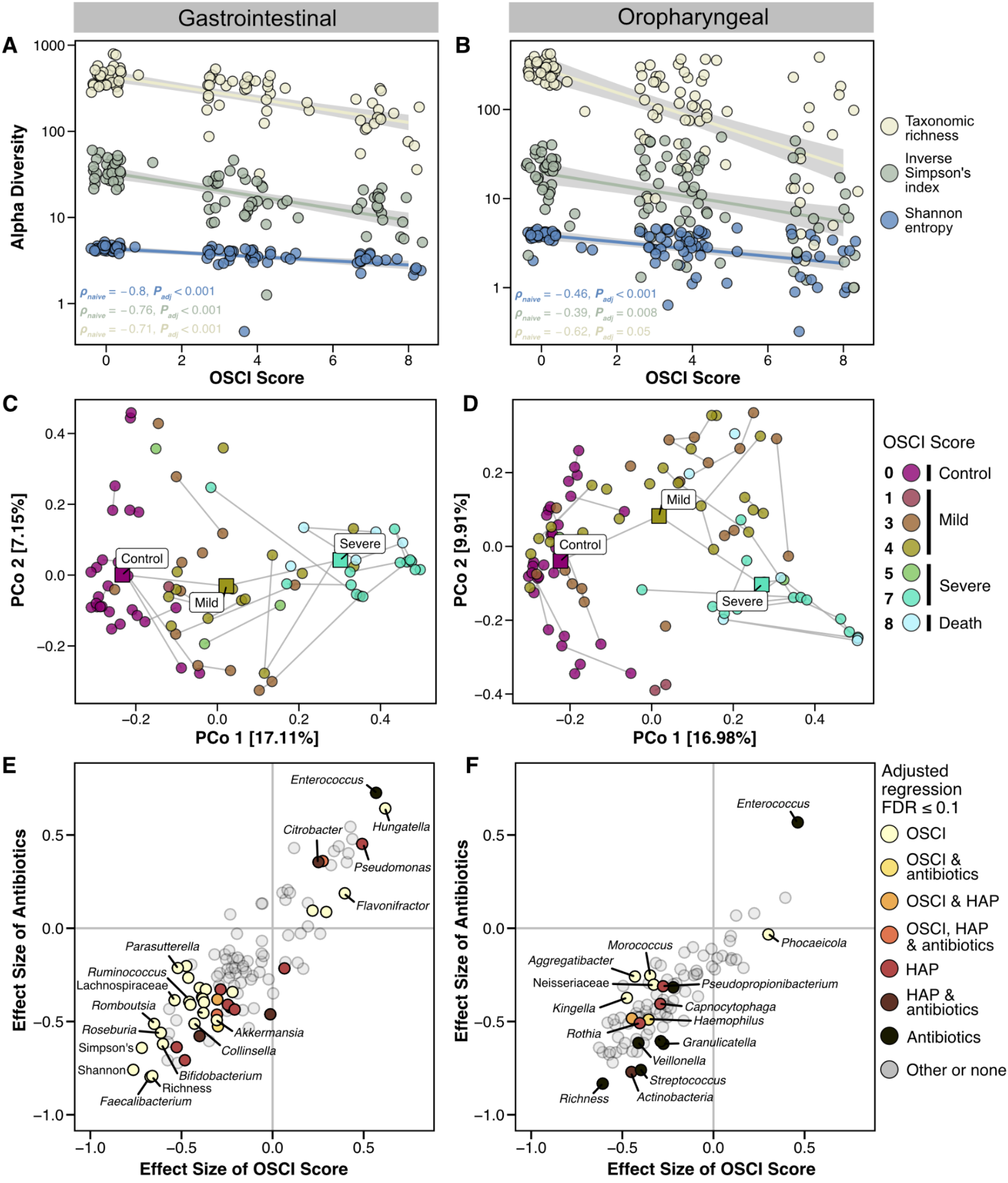
Microbiota compositional changes are associated with COVID-19 severity and hospitalization. **A-B)** Taxonomic richness and alpha diversity indices for gastrointestinal and oropharyngeal microbiota as a function of the worst disease severity attained by each individual (ordinal variable treated as continuous), with 0 indicating no infection (control group) and 8 indicating eventual death of the patient. Spearman correlations while controlling for antibiotic intake are shown. **C-D)** Principal coordinates analysis on mOTU (species) relative abundances with OSCI scores 1-4 indicating mild disease and 5-8 indicating severe or fatal disease. **E-F)** Standardized effect sizes for OSCI scores and current antibiotic use (Spearman correlation and Cliff’s delta, respectively) were calculated across all bacterial features in COVID-19 patients and are shown contrasted with one another. Results from our modeling of taxonomic and clinical data are overlaid, indicating the variables with the most explanatory power over a given taxonomic abundance (see Methods). See also Fig. S1 and S2. OSCI: ordinal scale for clinical improvement, HAP: hospital-acquired pneumonia.

In the oropharynx, severe disease courses were associated with depletion of e.g. *Aggregatibacter* and *Kingella*, while current antibiotic use was associated with decreased abundance of e.g. *Streptococcus* and enrichment of *Enterococcus* (Fig. 2F, Fig. S1). Similar to previous findings^30^, diversity and richness of the oropharyngeal microbiota as well as abundance of various members commonly found in healthy individuals are lower in COVID-19 patients as compared to uninfected controls, but these perturbations also appeared to be mainly driven by antibiotic administration (Fig. 2B, D, F, Fig. S1). In addition, we observed a negative correlation between the abundances of several commensals of the human oropharynx like *Actinobacteria* and *Rothia*, and the development of a secondary hospital-acquired pneumonia (HAP) (Fig. 2F). The depletion of *Actinobacteria* was at the same time partially explained by the use of antibiotics. Moreover, we found enrichment with Gram-negative bacilli like *Klebsiella* and *Achromobacter* in some of the TBS samples obtained from ventilated patients who developed HAP (for simplicity, we use the term HAP throughout to refer to all types of nosocomial pneumonia in both ventilated and non-ventilated patients) (Fig. S2). Collectively, our results indicate a direct association between intestinal microbiota abnormalities and severe COVID-19, whereas oropharyngeal microbiota disturbance appeared to be mainly driven by antibiotic use in our patients.

### Immune dysregulation in severe COVID-19

To characterize the systemic immune response in our cohort, we measured cytokines in plasma samples from all patients and healthy controls. In line with previous studies^29,30^, type I, II and III IFNs as well as several inflammatory cytokines including TNFα, IP-10/CXCL10, CCL2, and IL-10 were increased in early plasma samples of COVID-19 patients when compared to uninfected controls (Fig. 3A-H). While IFN levels mostly decreased in the later phase of the infection, production of the inflammatory cytokines remained high in severe COVID-19 patients. Next, we characterized PBMCs from a subset of patients at an early infection time point and uninfected controls (see Fig. 1) by droplet-based scRNAseq. Since we aimed to focus primarily on the innate immune cells, PBMCs were depleted of T and B lymphocytes before measurements. UMAP and cell type classification identified various cell types and subtypes expected in the mononuclear compartment of blood (Fig. 3I, J). Further analyses revealed an increase of classical monocytes in severe COVID-19 patients as compared to uninfected individuals and patients with mild infection (Fig. 3K), and a depletion of non-classical monocytes and cDCs in early COVID-19. NK cells were only depleted in patients with severe COVID-19. IFN-stimulated genes (ISG) were highly expressed in PBMCs of mild and severe COVID-19 patients (Fig. S3A,B,D), and ISG expression correlated with systemic levels of both type I and II IFNs (Fig S3C). Moreover, we measured expression of type I and III IFN genes in our oropharyngeal samples, and found increased *IFNL2* mRNA levels in mild COVID-19 patients as compared to uninfected controls (Fig. S3E). Overall, our results indicate that the systemic inflammatory response is disturbed in patients with severe COVID-19.

**Figure 3:**
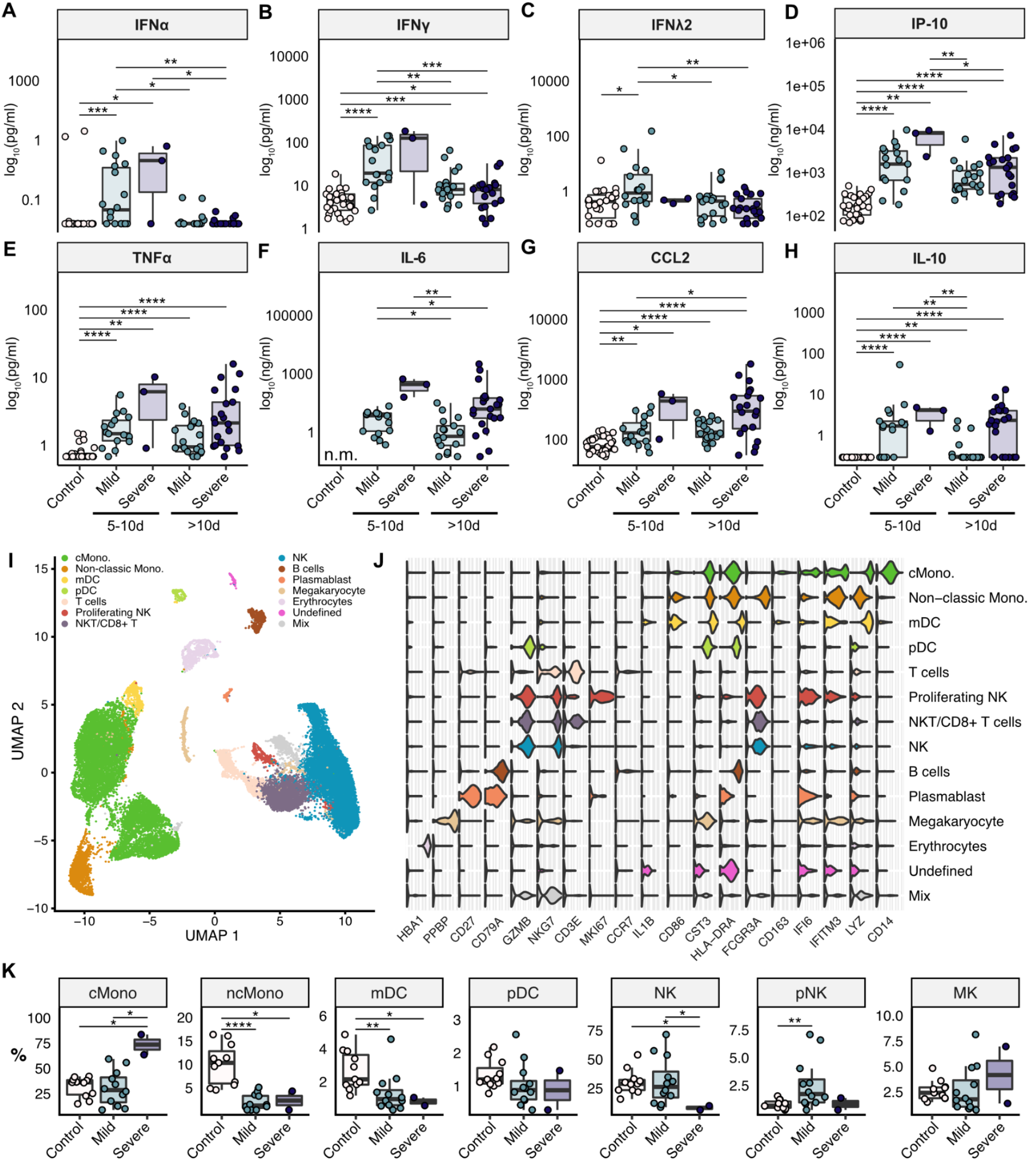
Dysregulated immune response in severe COVID-19 patients. **A-H)** Plasma levels of IFNs and inflammatory cytokines in healthy controls and COVID-19 patients were measured using MSD Meso Scale V-Plex assay kits or Simoa® Technology. **I)** UMAP representation of scRNA-seq profiles all merged T and B cell-depleted PBMC samples. 13 cell types were identified by cluster gene signatures. **J)** Violin plots showing top marker genes for the cell types shown in I). **K)** T and B cell-depleted PBMCs from 14 COVID-19 patients collected at an early infection phase (<10 days since symptom onset) and 11 healthy controls. The immune cell distribution varies between controls and COVID-19 patients and between mild and severe disease. Significant pairwise comparisons are denoted in panels A-H and K (Mann-Whitney U test). See also Fig. S3. n.m.: not measured; scRNAseq: single cell RNA sequencing, PBMCs: peripheral mononuclear blood cells, cMono: classical monocytes, ncMono: non-classical monocytes, mDC: myeloid dendritic cells, pDC: plasmacytoid dendritic cells, NK: natural killer cells, MK: megakaryocytes

### Alterations in tryptophan, bile acid, lipid, and amino acid metabolism in severe COVID-19

To identify potential microbiota- and/or host-derived factors underlying COVID-19 phenotypes, we performed metabolomic analyses on a total of 96 plasma and 77 urine samples from COVID-19 patients and uninfected controls. Our analysis highlighted significant differences in the plasma metabolome of COVID-19 patients when compared to uninfected controls, as well as direct associations between various metabolites and disease severity. In COVID-19 patients, we observed lower plasma levels of several tryptophan metabolites including the primarily dietary-derived tryptophan itself (*ρ*=-0.68, FDR-adjusted P<0.0001), the serotonin precursor 5-hydroxytryptophan (*ρ*=-0.38, FDR-adjusted P=0.0003), and the microbial metabolites tryptamine, indole-3-propionic acid, and indole-3-acetic acid^31^ (Fig. 4, Fig. S4A), indicating severe disturbance of host-dependent kynurenine and serotonin pathways and the microbiota-dependent indole metabolic pathway. Many of the altered tryptophan metabolites are ligands for the immunoregulatory aryl hydrocarbon receptor (AhR)^32^. In line with previous studies^33,34^, the host-derived tryptophan catabolites kynurenine, which is also an AhR ligand, and the potentially neurotoxic 3-hydroxykynurenine^35^ were strongly enriched in COVID-19 patients, with kynurenine showing a robust association with severity (*ρ*=0.7, FDR-adjusted P<0.0001; Fig. 4), in both early and late samples (Fig. S4A). Moreover, severe COVID-19 was robustly associated with lower plasma concentrations of the microbiota-produced secondary bile acid glycodeoxycholic acid (*ρ*=-0.42, FDR-adjusted P=0.0006), and with higher levels of the primary bile acid taurocholic acid (Fig. 4). Secondary bile acids have recently been shown to calibrate various immune cells and pathways^36–38^. In line with previous studies^39–41^, COVID-19 infection and disease severity were also associated with depletion of various lysophosphatidylcholines at all sampling timepoints and phosphatidylcholines in the early samples (Fig. S4A-B). Lysophosphatidylcholines are a group of proinflammatory lipids that are produced from phosphatidylcholine by the enzyme phospholipase A2, and shown to have effects on e.g. endothelial cells and immune cells^42,43^. Taken together, these results demonstrate that tryptophan, bile acid, lipid, and other amino acid metabolism is dysregulated in severe COVID-19.

**Figure 4:**
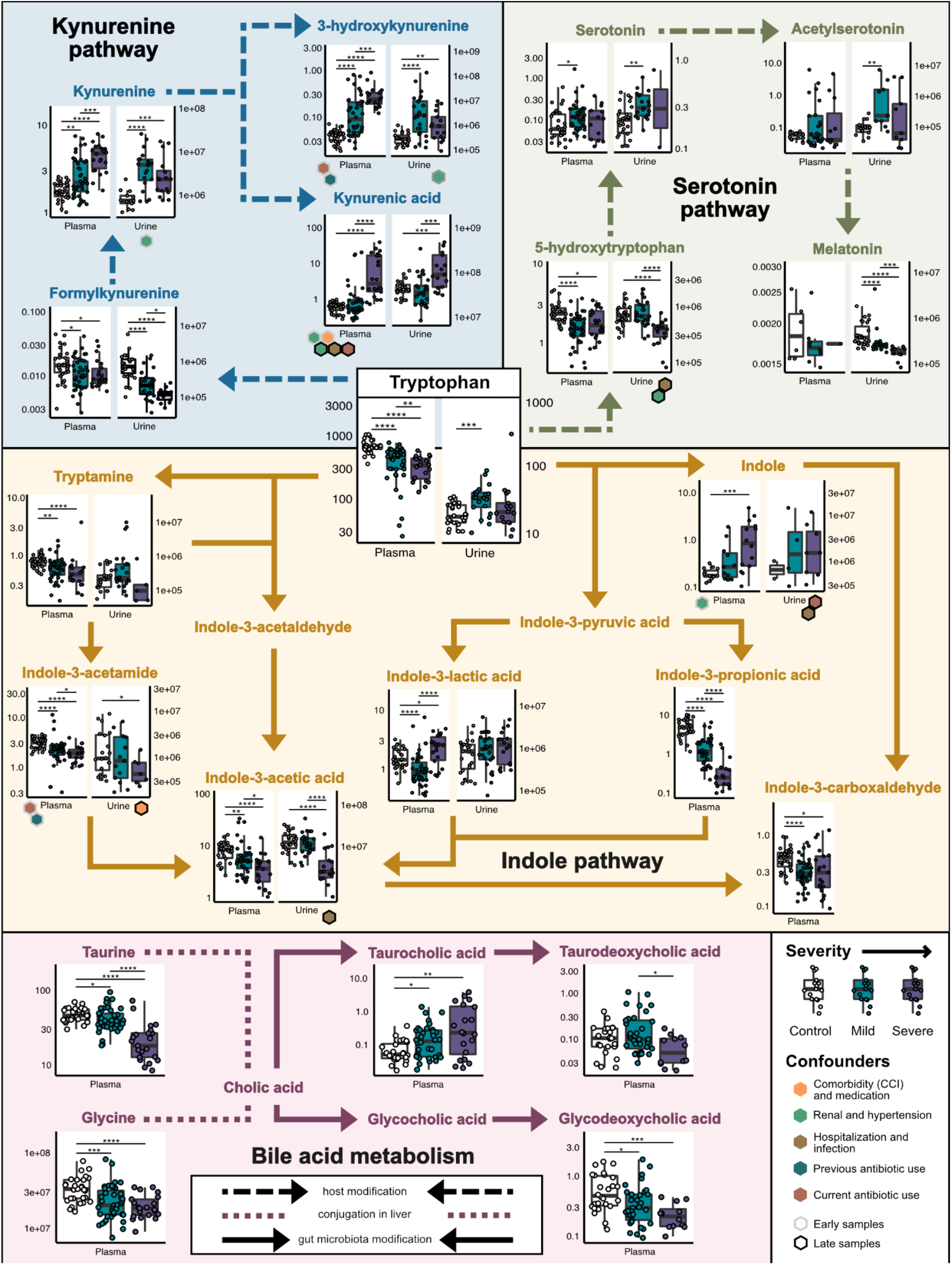
Tryptophan and bile acid metabolite associations with COVID-19 severity. Tryptophan and bile acid metabolite concentrations (given in ng/mL and μM, respectively) from all plasma and urine samples, annotated with pairwise Spearman test P-values and post-hoc identified confounders from early or late slices of the data. Some co-associated clinical variables were rationally grouped and relabeled here for annotation purposes, i.e. a metabolite confounded by hospitalization and infection reflects confounding by one or more of the following: HAP, number of days hospitalized, or other hospitalization-associated infections such as bacteremia and sepsis. The kynurenine and serotonin pathways are host-associated, while indole and part of the bile acid metabolism are carried out by gut microbes. See also Fig. S4. CCI: Charlson comorbidity index, HAP: hospital-acquired pneumonia

### Integrated analyses reveal associations between altered levels of tryptophan metabolites and enhanced production of proinflammatory cytokines

Finally, we integrated our various -omics data into our mixed-models analysis framework in order to establish potential associations between the microbiome, metabolome, and immune response parameters. First, we summarized the extent to which features from the different -omics spaces were associated with disease severity and/or confounded by different clinical factors such as previous or current antibiotic use, other medications, comorbidity, days of hospitalization, or age. This analysis revealed that many features of the gut microbiome, immune response, and plasma metabolome were robustly associated with COVID-19 severity, whereas most features of the oropharyngeal microbiome and urinary metabolome were only indirectly (i.e. confounded) or not significantly associated with COVID-19 severity, respectively (Fig. 5A, B). In particular, a large part of the alterations of the oropharyngeal microbiome appeared to be driven by previous and current antibiotic therapies (Fig. 5B). Next, we focused on the three most strongly COVID-19-linked -omics feature spaces, and used the robust severity-associated subset from each in further modeling steps to identify all unconfounded individual associations between features from different spaces (see Methods). This analysis uncovered 115 robust associations between the gut microbiome and the plasma metabolome, 3 between the gut microbiome and the immune response, and 29 between the plasma metabolome and the immune response (Fig. 5C). Lastly, we generated a model based on the most robust and interconnected associations between the three -omics data sets. Our model suggests that tryptophan metabolism is tightly linked to the dysregulated immune response in severe COVID-19, since a depletion of tryptophan and its metabolite 5-hydroxytryptophan, as well as enhanced levels of kynurenine and 3-hydroxykynurenine, explained significant increases in the production of IL-6, IFNγ and/or TNFα (Fig. 5D). Decreased 5-hydroxytryptophan in severe COVID-19 might be mediated by a depletion of Clostridiales and *Faecalibacterium* spp. from the gut, as both taxa robustly correlated with the metabolite levels, and in both cases the microbe-severity association was reducible to the microbe-metabolite association. Likewise, the association between IFNγ and disease severity was reducible to the association between IFNγ and kynurenine when modeled and tested jointly, indicating potential mediation. Depletion of *Faecalibacterium* was also associated with reduced production of the immunomodulatory metabolite tryptamine, which is similar to several of the above-mentioned tryptophan metabolites a ligand of AhR^32^. In addition, reduced intestinal abundance of Clostridiales explained significant variation in kynurenine, and in TNFα levels independent of 5-hydroxytryptophan. *Roseburia* might directly or indirectly be involved in the production of formylkynurenine, an intermediate of the kynurenine reaction and precursor for kynurenine and 3-hydroxykynurenine, since reduced abundances of this genus robustly associated with the decrease of the metabolite. Moreover, IFNγ production correlated with phenylalanine, and reduced concentrations of several lysophosphatidylcholines explained significant variation in IL-6, TNFα, and IFNγ levels. Taken together, our analysis indicates that alterations in both the microbiota- and host-dependent tryptophan metabolism, as well as potentially other metabolic pathways, may contribute to the dysregulated inflammatory immune reaction in severe COVID-19.

**Figure 5:**
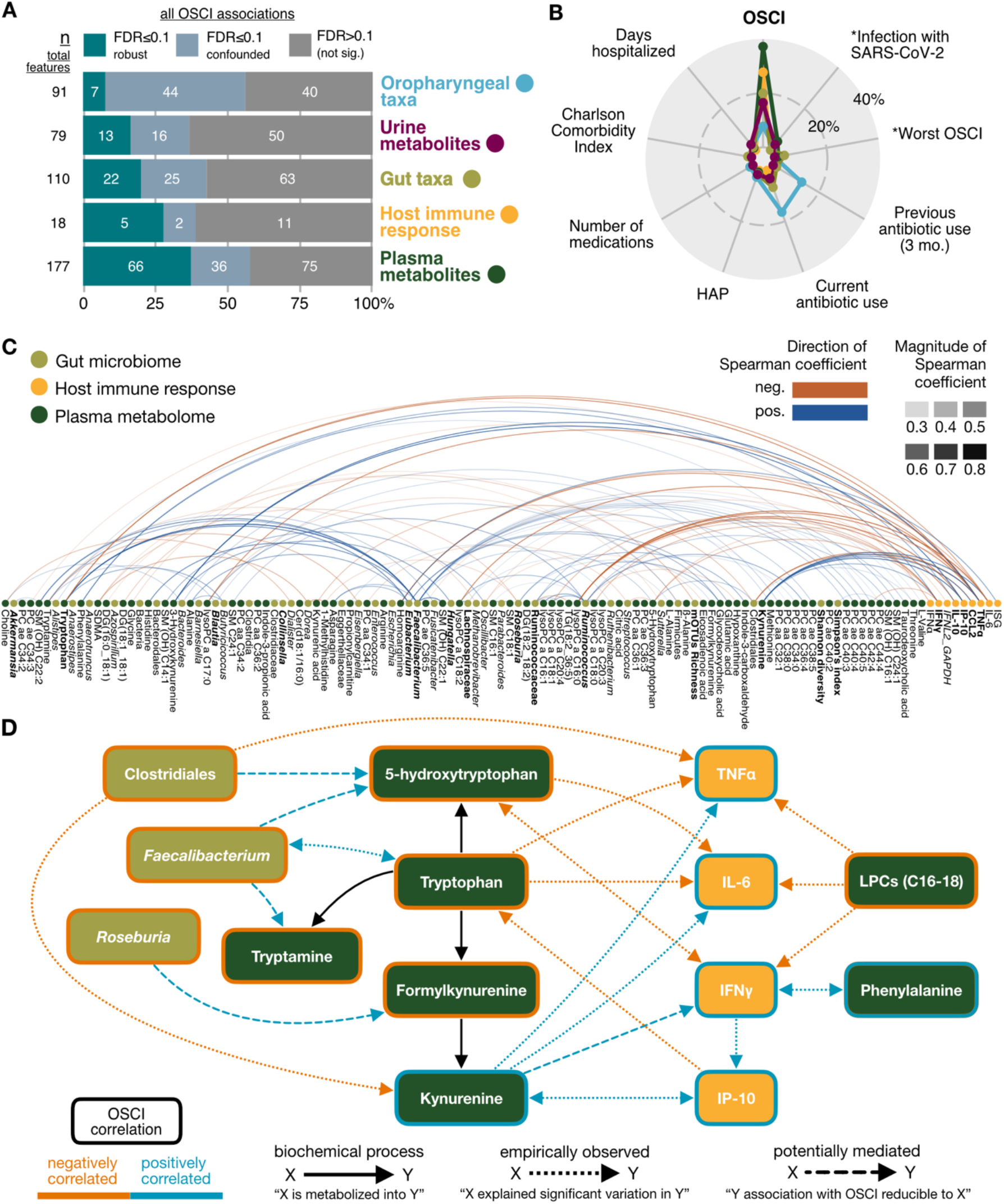
Integration of severity-associated models from among and across -omics spaces. **A)** Robustness of OSCI associations across all -omics features. Plasma metabolites were associated the most robustly with disease severity (highest n=66 and percentage of total features), while nearly half of the oropharyngeal taxa were confounded (n=44, 48%). **B)** Main confounding clinical variables. All significant OSCI associations are shown (cumulative area of non-grey bars from A), as well as an estimate of the percentage which were confounded, and if so by what. An asterisk* indicates co-association rather than confounding. **C)** Unconfounded associations between -omics spaces. Robust OSCI-associated features denoted in bold. **D)** Selected relationships across -omics spaces which emerged from our integrated linear model testing (see Methods). Some associations with disease severity were statistically reducible to other -omics features, which were considered potential mediators (rather than confounders) in this context. While low abundances of tryptophan and tryptophan-associated microbes negatively correlated with disease severity, downstream kynurenine metabolites were positively correlated with disease severity, and could explain significant variation in pro-inflammatory host cytokines (TNFα, IL-6) and type II interferon (IFNγ).

## Discussion

While several excellent microbiomics^20,24,27,44,45^, metabolomics^33^, and multi-omics studies^34,41,46–50^ of COVID-19 have been published, our work is unique in simultaneously measuring and analyzing a particularly large number of different -omics types, and, in this, the first to integrate gut and oropharyngeal metagenome sequencing, metabolomics, host transcriptomics, and cytokine profiling. Our analyses using linear mixed-effect models and exhaustive confounder testing revealed the plasma metabolome to be the -omics domain most affected by SARS-CoV-2 infection. Consistent with previous observations^33,40,41^, plasma levels of various host- and microbiota-derived tryptophan metabolites and lysophosphatidylcholines robustly correlated with COVID-19 severity, as did secondary bile acids in our study. In addition, enhanced inflammatory cytokine production and gut microbiota perturbations were strongly associated with the infection. For example, taxonomic diversity and richness were reduced in severe COVID-19 patients, and several potentially beneficial commensals (mainly belonging to the Clostridiales order as well as *Bifidobacterium*) were depleted, which is consistent with previous reports^20,21^. In contrast, the oropharyngeal microbiome alterations observed in our COVID-19 patients were largely explained by antibiotic use. Some of the changes (e.g. depletion of *Actinobacteria*) were at the same time associated with the development of HAP.

We related the depletion of intestinal Clostridiales and *Faecalibacterium* to the perturbed production of various tryptophan metabolites, and found that the increased production of key inflammatory cytokines including IL-6, TNFα, and IFNγ was statistically reducible to microbiota and metabolome features in some cases. The constellation of robust, cross-omics associations we uncovered thus contributes potential biomarkers as well as novel mechanistic hypotheses regarding the dysregulated immune response considered causative of severe COVID-19. In hospitalized patients, we estimated these mechanisms to further involve a vicious cycle, as critical illness, prolonged hospitalization, and high concentrations of inflammatory mediators further exacerbate the disruption of the microbiome and metabolome. Still, we speculate that several of our findings, e.g. hypotheses describing how certain intestinal commensals are associated with specific metabolites, or how tryptophan catabolites and lysophosphatidylcholines may regulate systemic cytokine production, are also relevant for other types of severe infections and perhaps non-infectious inflammatory diseases.

While levels of the host-derived tryptophan catabolite kynurenine were strongly elevated, tryptamine, indole-3-acetic and other microbiota-derived tryptophan catabolites were depleted in the plasma of severe COVID-19 patients. All of these metabolites are known to activate the AhR, which controls the differentiation and inflammatory potential of various innate and adaptive immune cells^31,51^. It is likely that, in aggregate, the markedly altered levels of these AhR ligands observed in our cohort contributed to the dysregulation of the immune response in severe COVID-19; however, further studies are needed to understand the cumulative impact of oppositely altered tryptophan catabolites (which presumably also differ with respect to their AhR binding affinities) on individual immune cells. Moreover, more research is also required to characterize the impact of these metabolites in different phases of COVID-19, such as the acute inflammatory phase, resolution, or the subsequent period of tissue repair.

We also observed that metabolites of the serotonin pathway were altered in plasma and/or urine of patients suffering from severe COVID-19, and that changes in 5-hydroxytryptophan levels explained significant variation in the production of IL-6 and IFNγ. From our integrated statistical modeling, we concluded that the depletion of 5-hydroxytryptophan was potentially mediated by low intestinal abundances of Clostridiales and *Faecalibacterium*, which is consistent with previous findings about the role of the microbiota in controlling the production of 5-hydroxytryptophan by colonic enterochromaffin cells^52^. Interestingly, 5-hydroxytryptophan has also recently been described to activate the AhR and to mediate CD8+ T cell exhaustion in antitumor immunity^53^. Thus, it appears reasonable to speculate that 5-hydroxytryptophan contributes to AhR-mediated calibration of inflammatory cytokine production during COVID-19, as well as to T cell exhaustion characteristic of severe SARS-CoV2 infection^54,55^.

Lysophosphatidylcholines are a group of bioactive lipids that have been found to influence e.g. immune and endothelial cells^43^. Low plasma levels of lysophosphatidylcholine have been associated with unfavorable outcomes in several chronic diseases^43^ and sepsis^56^. Moreover, lysophosphatidylcholine treatment was protective in mouse models of sepsis^57^. We observed reduced levels of lysophosphatidylcholines in severe COVID-19 patients, and strong associations with enhanced production of IL-6, TNFα, and IFNγ; this is potentially in line with a previous study showing that lysophosphatidylcholines inhibit the release of IL-6 by human monocytes in vitro^58^.

Another interesting group of metabolites whose production we found to vary with severe COVID-19 was the secondary bile acids. Secondary bile acids, which are converted from liver-derived primary bile acids by the microbiota, are known for their ability to influence various immune cells, e.g., via the receptors TGR5 and FXR^59^. Though we were not able to identify associations between secondary bile acids and specific immune features, this does not exclude the possibility that other immune mediators or cells in different compartments (such as the lung) are influenced by these metabolites. Indeed, a recent study uncovers how secondary acids control immunity against Chikungunya virus by enhancing type I IFN production by pDCs^36^. Further studies are required to evaluate the impact of secondary bile acids and other microbiota-derived metabolites on the immune response during COVID-19.

Our multi-omics study explores alterations in the microbiome, metabolome and immune response observed in severe COVID-19, and generates several novel testable hypotheses, but is not without limitations. First, we included only a relatively small number of patients from a single center, which mandates future validation in larger cohorts of patients; however, we performed deep phenotyping using various state-of-the-art omics techniques and integrative bioinformatics approaches, including extensive testing for confounding factors such as use of antibiotics and other medications, which are known to drive contradictory findings in the COVID-19 literature^60^. Second, for practical reasons and similar to probably all previously published work, we were unable to collect samples during the first days of infection, making it impossible to draw conclusions about mechanisms in the early phase of COVID-19. Third, we did not perform analyses of the immune response in the lung as the epicenter of the infectious event in COVID-19, which would be important for future studies. Overall, our study provides novel insights into the interaction between the microbiome, metabolome, and immune system in general, and into the pathogenesis of severe COVID-19 in particular. Future work should include animal experiments to mechanistically explore and resolve causality among the microbiome-metabolite-immune networks in COVID-19, and perhaps other infectious and inflammatory diseases. The disrupted microbiome-tryptophan metabolism-immune network described here might represent a potential target for novel intervention strategies to protect patients from severe COVID-19.

## Methods

### Study design and patient inclusion criteria

In the framework of the Pa-COVID-19, a prospective observational cohort study of patients with confirmed SARS-CoV-2 infection treated at Charité-Universitätsmedizin Berlin, we longitudinally collected stool, urine, TBS and blood samples as well as oropharyngeal swabs from hospitalized patients with COVID-19^26^. All patients with SARS-CoV-2 infection, as determined by positive PCR from respiratory specimens, who were willing to provide written informed consent were eligible for inclusion. Exclusion criteria included refusal to participate in the clinical study by patient or legal representative or clinical conditions that did not allow for blood sampling. The patients included in this study were enrolled between March 21 and June 15, 2020, before vaccinations or variants of concern. COVID-19 disease severity was classified to mild or severe disease according to the WHO clinical ordinal scale (https://www.who.int/publications/i/item/clinical-management-of-covid-19). Information on age, sex, medication, and comorbidities is listed in Table S1; unfortunately, information on diet and food intake while hospitalized was not recorded. Samples from uninfected individuals were collected in the framework of the COV-IMMUN study, a prospective study designed to analyze the immune response against SARS-CoV-2 and risk factors in health care workers at the Charité-Universitätsmedizin Berlin. The Pa-COVID-19 and COV-IMMUN studies are carried out according to the Declaration of Helsinki and were approved by the ethics committee of Charité-Universitätsmedizin Berlin (EA2/066/20, EA1/068/20). All patients or their legal representatives as well as the healthy individuals provided written informed consent for participation in the study.

### Metagenomic sample pre-processing, DNA extraction, and sequencing

Oropharyngeal swabs and stool samples were collected in collection tubes containing DNA/RNA shield (Zymo Research Cat# R1107-E and Cat# R1101) and frozen at -80C until further analysis was performed. DNA was isolated from the oropharyngeal swabs and stool samples using the ZymoBIOMICS™ DNA Miniprep Kit (Cat#D4300). For DNA isolation lysis of microbes was performed by mechanical disruption using a Mini-BeadBeater-96 (BioSpec) two times for 2 min. Libraries were prepared using the NEBNext Ultra II DNA Library Prep Kit (NEB Biolabs) according to manufacturer’s instructions. Sequencing was performed on the Illumina NovaSeq platform (PE150).

### PBMCs isolation and scRNA sequencing

PBMCs were isolated from heparinized whole blood by density centrifugation over Pancoll and cryopreserved in liquid nitrogen until further analysis. Frozen PBMC were recovered by rapidly thawing, and T and B cells were depleted by using CD19 and CD3 MicroBeads (Miltenyi Biotec Cat#130-097-055 and #130-097-043) to enrich for myeloid cells. Subsequently, the PBMC samples were hash-tagged with TotalSeq-A™ antibodies (Biolegend) and scRNA seq was performed by using a droplet-based single-cell platform (10xGenomics) as described recently^4^.

### ScRNAseq data analysis

The 10x Genomics CellRanger pipeline (v4.0.0) was used to pre-process the sequencing data. In brief, BCL files from each library were converted to FASTQ reads using bcl2fastq Conversion Software (Illumina) using the respective sample sheet with the 10x barcodes and TotalSeq antibodies utilized. Then, the reads were further aligned to the reference genome provided by 10x Genomics (Human reference dataset refdata-cellranger-GRCh38-3.0.0) and a digital gene expression matrix was generated to record the number of UMIs for each gene in each cell. Next, the expression matrix from each library was loaded to R/Seurat packages^61^ (v4.0.1) for downstream analysis. To control the data quality, we further excluded low-quality cells with >15% mitochondrial reads, < 100 or > 3,000 expressed genes, or < 500 UMI counts. In addition, genes expressed in less than three cells were also excluded from further analysis. After QC, we normalized the gene counts from each cell, where original gene counts were divided by total UMI counts, multiplied by 10,000 (TP10K), and then log-transformed by log10(TP10k+1). We then scaled the data, regressing for total UMI counts, and performed principal component analysis (PCA) based on the 2,000 most-variable features identified using the vst method implemented in Seurat. Cells were then clustered using the Louvain algorithm based on the first 20 PC dimensions with a resolution of 0.3. For visualization, we applied UMAP based on the first 20 PC dimensions. The obtained clusters were annotated by the expression of PBMC marker genes. The expression of selected genes was visualized by violin plots.

### Quantitative reverse transcription PCR

For measuring expression of type I and II IFNs in the upper airways, total RNA was isolated from oropharyngeal swab fluid using mirVana™ miRNA Isolation Kit (Cat# AM1561). The RNA was reverse-transcribed using the high capacity reverse transcription kit (Applied Biosystems, Darmstadt, Germany), and quantitative PCR was performed using TaqMan assays (*GAPDH*: Hs02786624_g1, *IFNL2*: Hs04193048_gH, *IFNB*: Hs01077958_s1 Life Technologies, Darmstadt, Germany) on an ABI 7300 instrument (Applied Biosystems, Darmstadt, Germany). The input was normalized to the average expression of GAPDH and relative expression (relative quantity, RQ) of the respective gene in the healthy control individuals was set as 1.

### Cytokine ELISA

Plasma concentrations of IL-10, IL-12p70, IL-17A, IL-1α, IL-1β, IL-4, IL-22, IP-10, MCP-1, TNFα, IFNγ were measured by using MSD Meso Scale V-Plex assay kits (Meso Scale Diagnostics). Plasma concentrations of IFNα and IL-28A were quantified by using Simoa® Technology (Quanterix Corporation).

### Viral load measurements

SARS-CoV-2 RNA detection and quantification in respiratory swabs and stool samples was done as described before^62,63^ and by using either the cobas® SARS-CoV-2 test on the cobas® 6800/ 8800 system or the SARS-CoV-2 E-gene assay from TibMolbiol on a Roche MagNApure 96/LightCycler 480er workflow. Viral RNA concentrations were calculated by using the CT-Value of the E-gen target and by applying calibration curves of quantified reference samples and in-vitro transcribed RNA^64,65^.

### Plasma and urine metabolomic sample pre-processing

Plasma and urine samples were prepared in four different ways depending on the metabolite of interest. Urine samples were treated with urease before Biocrates and CCM analyses. For measuring SCFA, tert-Butyl-Methyl-Ether (MTBE; Sigma 650560) HCl (37%; Roth X942.1) and crotonic acid as internal standard (Sigma 113018-500G) were added to the plasma and urine samples. Unfortunately, there were technical issues with the analytical measurements resulting in the SCFA measurements being excluded from further analysis on quality control grounds. A broad metabolite analysis was conducted using a Biocrates MxP Quant 500 kit. For safety reasons, samples were measured after adding 100% ethanol (LC-MS grade; Fisher Scientific) to the plasma and urine samples. For the analysis of tryptophan derivatives, an extraction solvent (89,9% Methanol in 0,2% FA and 0,02% ascorbic acid) was added to the plasma and urine samples. The preparation of the plasma and urine samples for CCM GC-MS analysis consisted of adding 100% Methanol (LC-MS grade; Fisher Scientific).

### Biocrates MxP Quant 500 assay and measurement

Plasma or urine was added in a 1:2 dilution to ethanol (EtOH, Fisher Scientific, New Hampshire, US; 50 μL to 100 μL EtOH) and vortexed for 20 seconds. Samples were stored at -80°C until use. The MxP Quant 500 kit from Biocrates Life Science AG is a fully automated assay based on phenylisothiocyanate (PITC) derivatization of the target analytes using internal standards for quantitation. Plate preparation was done according to the manufacturer’s protocol. Briefly, 30 μL of the diluted plasma or urine was transferred to the upper 96-well plate and dried under a nitrogen stream. Thereafter, 50 μL of a 5% PITC solution was added. After incubation, the filter spots were dried again before the metabolites were extracted using 5 mM ammonium acetate in methanol (MeOH, Fisher Scientific, New Hampshire, US) into the lower 96-well plate for analysis after further dilution using the MS running solvent A. Quality control (QC) samples were prepared by pooling plasma or urine from each sample.

Evaluation of the instrument performance prior to sample analysis was assessed by the system suitability test (SST) according to the manufacturer’s protocol. The LC-MS system consisted of a 1290 Infinity UHPLC-system (Agilent, Santa Clara, CA, USA) coupled to a QTrap 5500 (AB Sciex Germany GmbH, Darmstadt, Germany) with a TurboV source. Acquisition method parameters and UHPLC gradient for LC and FIA mode are shown in Table S9-11. All compounds were identified and quantified using isotopically-labeled internal standards and multiple reaction monitoring (MRM) for LC and full MS for FIA as optimized and raw data was computed in Met*IDQ*™ version Oxygen (Biocrates Life Science AG, Innsbruck, Austria). A script developed in-house (MetaQUAC) was used for data quality analysis and preprocessing^66^. Quality assurance and control were reported using the recommended standards by mQACC (Table S8).

### Gas chromatography mass spectrometry (GC-MS) measurement of key central carbon pathway metabolites

MeOH containing 2 μg/mL cinnamic acid as internal standard (Sigma Aldrich, St. Louis, Missouri, US) was aliquoted (112.5 μL) and stored on ice. 25 μL of plasma was added to the MeOH followed by addition of 329 μL MeOH, 658 μL chloroform (CHCl_3_, Sigma Aldrich, St. Louis, Missouri, US), and 382.5 μL water (H_2_O, Fisher Scientific, New Hampshire, US). Samples were vortexed and left on ice for 10 minutes to separate into a biphasic mixture. The samples were centrifuged at 2,560xg for 20 minutes at 4°C and then left to equilibrate at room temperature for 20 minutes. 300 μL of the upper polar phase was then collected and dried in a rotational vacuum concentrator (Martin Christ, Osterode, Germany). To the urine samples (150 μL), 200 μL of 1 mg/mL urease solution in water was added, sonificated for 15 minutes and left on ice for 45 minutes. Ice cold MeOH (800 μL containing 2 μg/mL cinnamic acid as internal standard) was added, vortexed and centrifuged at maximum speed for 10 minutes at 4°C. The supernatant (750 μL) was transferred to a new vial and stored at -80°C until use. Urine samples were normalized to the according osmolarity and dried in a rotational vacuum concentrator (Martin Christ, Osterode, Germany). Quality control (QC) samples were prepared by pooling the extracts of plasma or urine from each sample.

For derivatization the extracts were removed from the freezer and dried in a rotational vacuum concentrator (Martin Christ, Osterode, Germany) for 60 min before further processing to ensure there was no residual water which may influence the derivatization efficiency. The dried extracts were dissolved in 15 μL or 20 μL of methoxyamine hydrochloride solution (40 mg/mL in pyridine, both Sigma Aldrich, St. Louis, Missouri, U) and incubated for 90 min at 30 °C with constant shaking, followed by the addition of 50 μL or 80 μL of N-methyl-N-[trimethylsilyl]trifluoroacetamide (MSTFA, Macherey-Nagel, Düren, Germany) and incubated at 37 °C for 60 min for plasma and urine, respectively. The extracts were centrifuged for 10 min at 18,213 xg, and aliquots of 25 μL (plasma) or 30 μL (urine) were transferred into glass vials for GC-MS measurements. QC samples were prepared in the same way. An identification mixture for reliable compound identification was prepared and derivatized in the same way, and an alkane mixture for a reliable retention index calculation was included (10.3390/metabo10110457). The metabolite analysis was performed on a Pegasus 4D GCxGC TOFMS-System (LECO Corporation) complemented with an auto-sampler (Gerstel MPS DualHead with CAS4 injector). The samples were injected in split mode (split 1:5, injection volume 1 μL) in a temperature-controlled injector with a baffled glass liner (Gerstel). The following temperature program was applied during the sample injection: for 2 min, the column was allowed to equilibrate at 68 °C, then the temperature was increased by 5 °C/min until 120 °C, then by 7 °C/min up to 200 °C, then by 12 °C/min up to a maximum temperature of 320 °C, which was then held for 7.5 min. The gas chromatographic separation was performed on an Agilent 7890 (Agilent Technologies), equipped with a VF-5 ms column (Agilent Technologies) of 30 m length, 250 μm inner diameter and 0.25 μm film thickness. Helium was used as the carrier gas with a flow rate of 1.2 mL/min. The spectra were recorded in a mass range of 60 to 600 m/z with 10 spectra/second. Each sample was measured twice (technical replicates). The GC-MS chromatograms were processed with the ChromaTOF software (LECO Corporation) including baseline assessment, peak picking and computation of the area and height of peaks without a calibration by using an in-house created reference and library containing the top 3 masses by intensity for 42 metabolites (55 intermediates; Table S12) related to the central carbon metabolism.

The data were exported and merged using an in-house written R script. The peak area of each metabolite was calculated by normalization to the internal standard cinnamic acid. Relative quantities were used. CCM and tryptophan data were batch corrected using the cubic spline drift correction from notame^67^ (v0.0.5, in R v4.0.1) followed by QC-sample median normalization^68^. Urine tryptophan data was only QC-sample median normalized. Quality assurance and control were reported using the recommended standards by mQACC (Table S8).

### Tryptophan metabolite analysis using UPLC-MS

For the tryptophan analysis, liquid chromtography – mass spectrometry (LC-MS) analysis was performed with a 1290 Infinity 2D HPLC system (Agilent Technologies, USA) combined with a TSQ Quantiva triple quadrupole mass spectrometer with a heated ESI source (Thermo Scientific, USA). Before starting, an extracting solvent was prepared comprising 90% methanol, 0.15 μg/mL mixed internal standards, 0.02% ascorbic acid, and 0.2% formic acid. This was placed at -20°C to cool. For urine samples a 1:5 (v/v) dilution was prepared in water prior to a urease digestion at 37°C for 40min with 10U urease (Sigma Aldrich). For each sample, 280 μL pre-chilled extracting solvent was added to 140ul of plasma or urease digested urine. Samples were held at 4°C and shaken for 10 min at 1000rpm (Eppendorf ThermoMixer C), incubated at -20°C before being centrifuged for 15 min at 11000g and 4 oC. The supernatant was transferred to a dark LC-MS vial for LC-MS/MS analysis. 20μl of each plasma sample was pooled, and the pooled plasma was also extracted to make quality control (QC) samples. These QC samples were run every 6 samples. LC-MS analysis of 5μl injection was combined with a triple quadrupole mass spectrometer using a 10-min gradient. A reversed-phase column was used (VisionHT C18 Basic; L × I.D. 150 mm × 4.6 mm, 3 μm particle size, Dr Maisch, Germany) and held at a constant temperature of 30°C. The mobile phase consisted of 0.2 % formic acid in H2O (solvent A) and 0.2 % formic acid in methanol (solvent B). The following gradient was run with a constant flow rate of 0.4 ml min−1: A/B 97/3 (0 min), 70/30 (from 1.2 min), 40/60 (from 2.7 to 3.75 min), 5/95 (from 4.5 to 6.6 min) and 97/3 (from 6.75 to 10 min). The molecular ion and at least two transitions were monitored for the 15 metabolites that are part of the tryptophan pathway.

Data was exported into Skyline (v.19.1, 64-bit) to identify and quantify peak intensity and area. Transition settings in the Skyline search were: isotopic peaks included: count; precursor mass analyzer: QIT; acquisition method: targeted; product mass analyzer: QIT. Method match searching tolerance was 0.6 m/z, and data was manually checked to ensure the correct peaks were selected. Cubic spline drift correction was applied per metabolite and to all sample types using the pooled quality control (QC) samples as references to fit the splines. The first and the last QC samples used to fit the cubic splines are the most critical to the resulting fit. In this case, the last conditioning pooled QC sample and the first of two pooled QC replicates at the end of the analytical run were used as the first and last QC respectively in the batch correction. Standard samples of increasing concentration were used to construct calibration curves using linear fits per metabolite in ng/ml. Concentration values less or equal to zero were declared as missing. In this study, several study groups featured measurements systematically outside of the calibration range for some metabolites (i.e. below or above the smallest or largest standard sample applied for a metabolites calibration curve, respectively). Further, calibration of QC standard samples resulted in insufficient accuracy. However, precision (%RSD in either standard or pooled QC samples) was adequate for most compounds. Hence, metabolites cannot be considered as absolutely but as relatively quantified in this study. In-house R scripts were used for internal standard normalization, calibration, statistics and plotting. Quality assurance and control were reported using the recommended standards be mQACC (Table S8):

### Taxonomic and functional microbiome profiling

Whole genome shotgun sequencing reads were analyzed using the NGLess pipeline (v1.3.0). Sequences were quality controlled, trimmed and merged using NGLess defaults and subsequently filtered for human reads. One stool sample and two TBS samples were of insufficient quality and removed from further analysis. Taxonomic assignment was performed using the mOTUs profiler (v2.6). Functional annotation was performed by mapping rarefied reads to the Global Microbial Gene Catalog (GMGC, v1.0), then binning into either KEGG Modules through custom scripts or gut metabolic modules (GMMs) using the Java implementation of the Omixer-RPM reference mapper software.

### Taxonomic profile post-processing, normalization, and diversity analyses

Both mOTUs and functionally-profiled metagenomic counts were transformed to relative abundances, then filtered to exclude features which were a) nonzero in less than 10% of samples, b) with a mean relative abundance less than 10e-4, or c) with zero variance. For most analyses with the shotgun metagenomic data, as stated where results are referenced, manually binned genus-level mOTU counts were used to increase the strength of the signal, which the integration analysis included an additional filtering step to further refine (more detail in that section, below). The *vegan* (v2.5.7) and *stats* packages were used for alpha and beta diversity calculation. The beta diversity principal coordinates analysis used the (species-level) mOTU counts.

### Statistical testing of -omics data and post-hoc confounder analysis with clinical variables

All statistical analysis was performed with R (v4.0.3) using the *targets* workflow manager (v0.12.1) and *renv* environment manager (v0.12.5) to enhance reproducibility. All figures were generated using *ggplot2* (v3.3.3) and *patchwork* (v1.1.1). Testing was performed using the *metadeconfoundR* package (v0.2.7) as described in Forslund *et al*.^69^ (especially Extended Data Figure 1 for a graphical overview) and briefly described here.

To identify which clinical variables associated with -omics features, standardized, non-parametric effect sizes (Cliff’s delta and the Spearman correlation for binary and continuous variables, respectively) were calculated and tested for significance in a first step. The full set of clinical variables and per-individual values are given in Table S1; variables which had less than three nonzero observations in both the patient and control groups (6 total) were not tested.

In a second step, significant clinical variables from the first step were used in an iterative, nested regression procedure to assess post-hoc confounding potential. Single -omics feature abundances were rank-transformed and regressed onto a disease status label, both 1) with and 2) without a potentially confounding variable from the first step, followed by a likelihood ratio test (LRT) between nested models 1 and 2. Linear mixed-effect models were used to account for longitudinal sampling. This was repeated combinatorially across all post-processed -omics feature abundances and clinical variables, and integrated to yield a single status for each feature-clinical variable association (including disease status and severity): robust (not confounded by any naively significant covariates), confounded (and if so by what/which), or not significant (summary of results shown in Fig. 5A). The Benjamini-Hochberg procedure was used to correct for multiple testing in both the naive statistical tests and likelihood ratio tests.

Concretely, our software produces a table of results in which each row contains the statistical summary for a single -omics feature and clinical variable pair (e.g. alpha diversity and OSCI score, respectively). There are identifier columns for each of these (“Y_dep.var” and “X_ind.var” for dependent Y and independent X variables in the models, respectively), and columns for naive effect size and adjusted p-values (XY_eff.size and XY_p.adj, respectively). The final column (“AssocStatus”) is either a status (D for “deconfounded” or NS for “not significant”), or, if “confounded”, a list of other clinical variables which resulted in a no-longer-significant association between X and Y in the given row when modeled as second independent variable. In our example, the gut alpha diversity was “deconfounded” (i.e. robustly associated with the OSCI score), while the oropharyngeal alpha diversity was confounded by antibiotic use, which was listed (see Fig. S1). Our combined results from analysis with each of the post-processed -omics data tables and clinical metadata are given in Table S6.

### Cross-omics associations and integrated statistical analysis

As described above, all individual -omics features were tested for associations with the same set of clinical factors including disease status and severity, revealing a subset of features from each space which was robustly correlated with the OSCI score (Fig. 5A, B). To examine associations and generate hypotheses between different -omics spaces, we reconfigured our statistical framework to include robust subsets of severity-associated “cross-omics” features as additional independent variables. This produced naive correlations between e.g. severity-associated microbial taxa and metabolites or immune parameters, and further expanded our ability to classify their robustness via iterative nested model testing.

As a concrete example of a single step in this extended framework: plasma kynurenine (KYN) and IFNγ were both robustly associated with severity, so kynurenine was included as an additional independent variable when re-testing IFNγ against disease severity (OSCI). Three models were built:

osci_model: rank(IFNγ) ∼ OSCI

kyn_model: rank(IFNγ) ∼ KYN

full_model: rank(IFNγ) ∼ OSCI + KYN

Then two likelihood ratio tests (LRTs) were performed with different nested model comparisons, and their results were used to classify the association between IFNγ and the OSCI score:

Test 1: likelihood ratio test between full_model and osci_model

Test 2: likelihood ratio test between full_model and kyn_model

Test 1 checks whether kynurenine explains significant variation in IFNγ measurements beyond that which is already explained by the OSCI score, while test 2 checks the converse. If only test 1 is significant, then, it can be concluded that the OSCI-IFNγ association is statistically reducible to the KYN-IFNγ association, and therefore the OSCI-IFNγ association may be considered “confounded” by kynurenine. If only test 2 or both tests are significant, the OSCI-IFNγ association is at least partially statistically independent of kynurenine, and may be considered robust (so long as it remains statistically independent from the other clinical and cross-omics variables tested).

This logic is identical to that used in the framework to test each set of -omics features against clinical variables; however, the aim there was to reduce dimensionality in each -omics space to a robust severity-associated subset. Here, we broadened the interpretation of post-hoc LRT-identified “confounders” in this cross-omics context to consider them as potential mediators. Our combined results needed to generate Fig. 5C and 5D are given in Table S7.

## Data and Code Availability

Raw sequencing data will be deposited and made publicly available before publication. The metabolomics data are available on MetaboLights^70^ with the unique identifier MTBLS6600 (www.ebi.ac.uk/metabolights/MTBLS6600). All supplemental, processed data tables are uploaded separately and the code to perform the confounder and integrated statistical analyses are hosted at https://github.com/sxmorgan/pa-covid-multi-omics. Any further information required to reanalyze the data in this manuscript is available from the lead author upon request

## Acknowledgments

The authors are grateful to all patients, their relatives, and volunteer controls for consenting to biosampling and data collection. We thank all members of the Pa-COVID collaborative study group. We thank Alina Eisenberger (BIH Metabolomics Platform) for technical assistance. This work was supported by start-up funding by the Berlin Institute of Health (BIH) to JAK, SKF, and BO, and by the Deutsche Forschungsgemeinschaft (SFB-TR84 A1/A5) to BO. TRL and TS were supported by the VolkswagenStiftung’s initiative “Niedersächsisches Vorab and Ministry of Lower Saxony (MWK, Project 76251-99). SKF was supported by the German Center for Cardiovascular Research, the German Research Council (projects SFB1365, SFB1470 and KFO339) and the German Ministry of Education and Research. EW and ML are supported by the Project “Virological and immunological determinants of COVID-19 pathogenesis – lessons to get prepared for future pandemics (KA1-Co-02 ‘COVIPA’)”, a grant from the Helmholtz Association’s Initiative and Networking Fund. CT was supported by the Deutsche Forschungsgemeinschaft (project number 400667201). VC was supported by the German Ministry of Education and Research through project VARIPath (01KI2021). VC is a participant in the BIH–Charité Clinician Scientist Program funded by Charité—Universitätsmedizin Berlin and the Berlin Institute of Health.

## Author Contributions

BO and SKF conceived and supervised the work with contributions from JAK. BMPL, MK, FK, LM, NS, CT and LES designed the clinical study and organized sample collection and processing. BMPL prepared the stool samples and oropharyngeal swabs with help from SC, which AAB, TRL and TS sequenced, and UL processed. BMPL prepared the plasma and urine samples with help from FFV and IR, and UB, RFG, and MK processed the metabolite data. BMPL isolated the PBMCs with help from FFV and SB, which EW and ML sequenced and processed. BZ and YL analyzed the single cell data. VC assessed viral loads. AM and CM performed multiplex cytokine assays. ME integrated all processed data, and conceived and performed the statistical analyses. ME designed and produced the figures with contributions from BMPL, BO, UB and BZ. BMPL, BO, ME and SKF developed the hypotheses and interpreted the results with input from JAK. BO, ME, and BMPL wrote the original draft of the manuscript with input from SKF. All authors discussed and approved the final manuscript.

