## Supplementary Information for "Gut microbiota dysbiosis is associated with altered tryptophan metabolism and dysregulated inflammatory response in severe COVID-19"

Supplemental Information


Figure S1: Severity-associated microbiota in hospitalized COVID-19 patients and controls, related to Figure 2

Figure S2: Dominant respiratory taxa in hospital-acquired pneumonia (HAP) patients, related to Figure 2

Figure S3: IFN response in COVID-19 patients, related to Figure 3

Figure S4: Metabolite associations with disease severity at both early and late timepoints, related to Figure 4

Table S1: Cohort characteristics, related to Table 1

Table S2: Clinical lab parameters, related to Figure 1

Table S3: Taxonomic profiles, related to Figures 2, S2

Table S4: Immune response profiles, related to Figures 3, S3

Table S5: Metabolite profiles, related to Figures 4, S4

Table S6: Results from statistical analysis with clinical variables, related to Figures 2-4, S2-S4

Table S7: Integrated statistical analysis results, related to Figure 5

Table S8: Quality Control reporting of metabolomics data according to mQACC standards

Table S9: Ultra-high performance liquid chromatography (UHPLC) gradient for LC part

Table S10: Ultra-high performance liquid chromatography (UHPLC) gradient for flow injection (FIA) part

Table S11: Acquisition method parameters for liquid chromatography (LC) and flow injection analysis (FIA)

Table S12: List of metabolite derivatives and their biological groups used for reference search

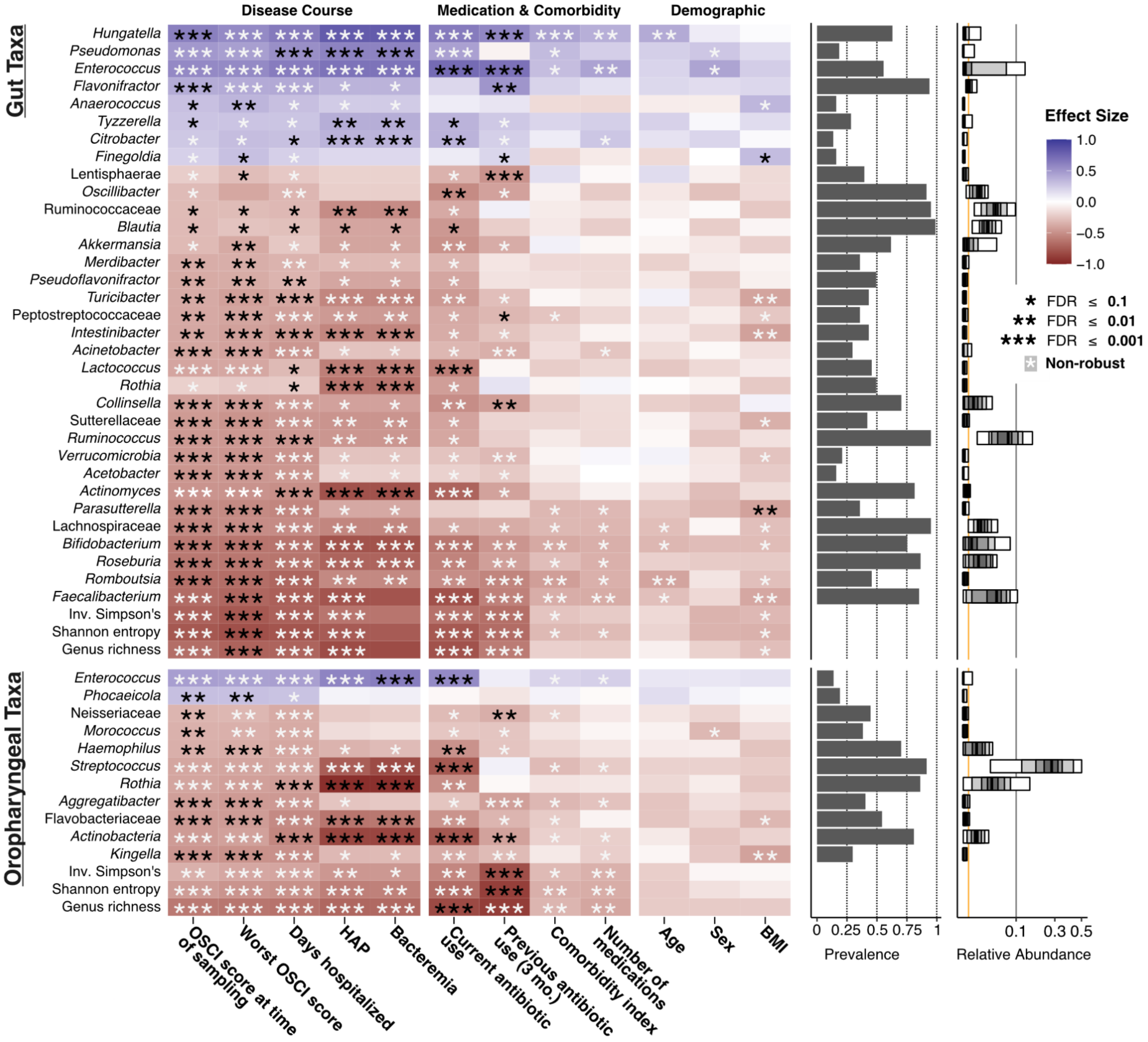


**Figure S1: Severity-associated microbiota in hospitalized COVID-19 patients and controls, related to Figure 2**
To correlate bacterial taxa with clinical variables (see Table S1 for full list), standardized, non-parametric effect sizes were calculated and tested for significance (shown in the heatmap, either Spearman for continuous variables or Cliff's delta/MWU for binary variables). Summary alpha diversity metrics and (naïve) severity-associated taxa of the gut and oropharynx are depicted. In a second step, feature-wise iterative models were built and integrated to determine the robustness of clinical associations, including disease severity, and identify potential confounders (see Methods). If a naïve association retained significance across all possible covariate models, its significance is denoted in black in the heatmap and it may be considered robust. White stars denote instances in which naive associations were reducible to one or more other covariates and therefore are not considered robust. Multiple columns of robust associations for a given feature indicate variables which captured similar variation, e.g. hospitalization-associated variables (length of stay, HAP, bacteremia). Taxa prevalences and square root-transformed relative abundance quantile plots are shown on the right; the yellow line indicates a relative abundance of 0.001, which was used as a cutoff to determine a robust OSCI-associated subset of microbiota features for later integration with host metabolites and immune response features. See also main text Fig. 1. HAP: hospital-acquired pneumonia; OSCI: ordinal scale for clinical improvement

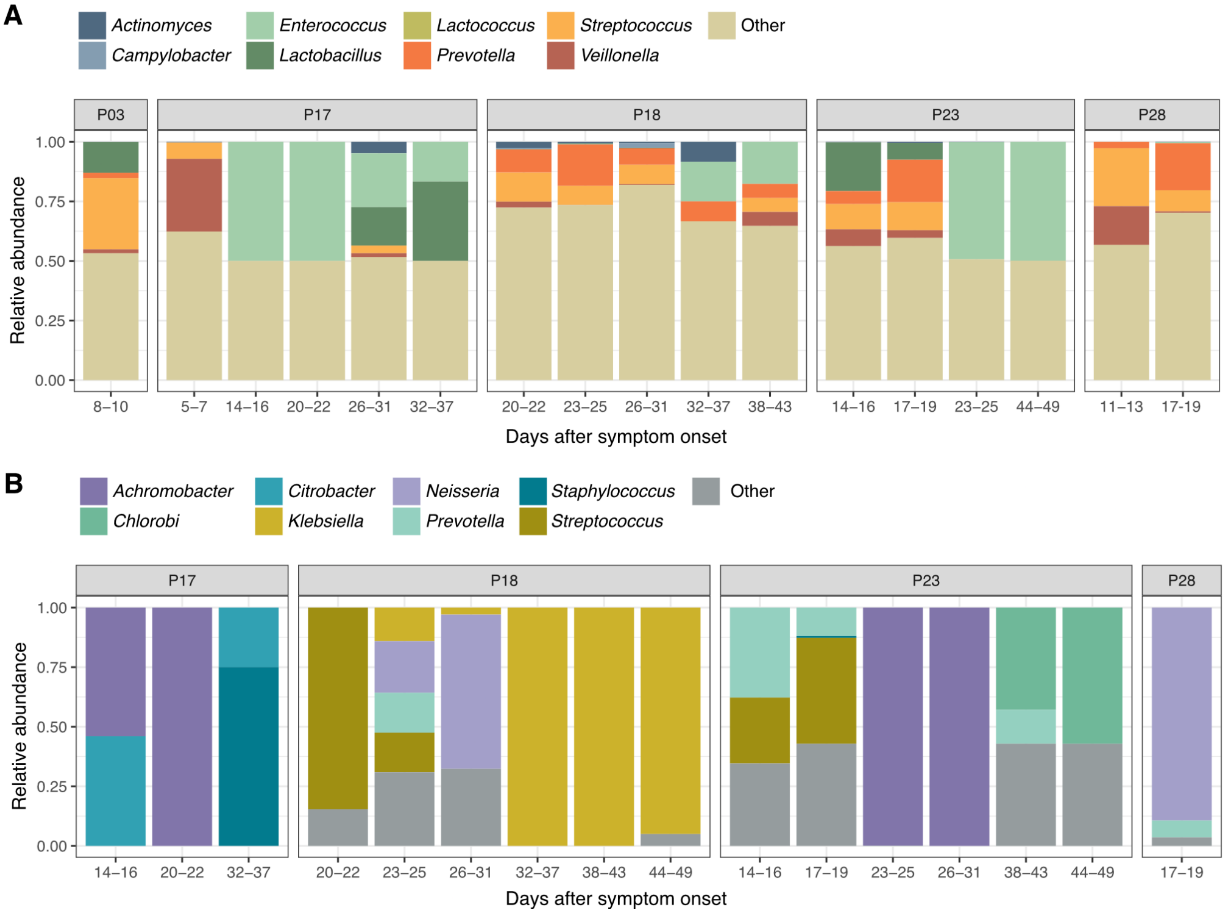


**Figure S2: Dominant respiratory taxa in hospital-acquired pneumonia (HAP) patients, related to Figure 2**
**A)** Oropharyngeal samples from the five patients with nosocomial pneumonia infections; all were ventilated except for P03. **B)** Tracheobronchial sputum samples obtained from ventilated patients.

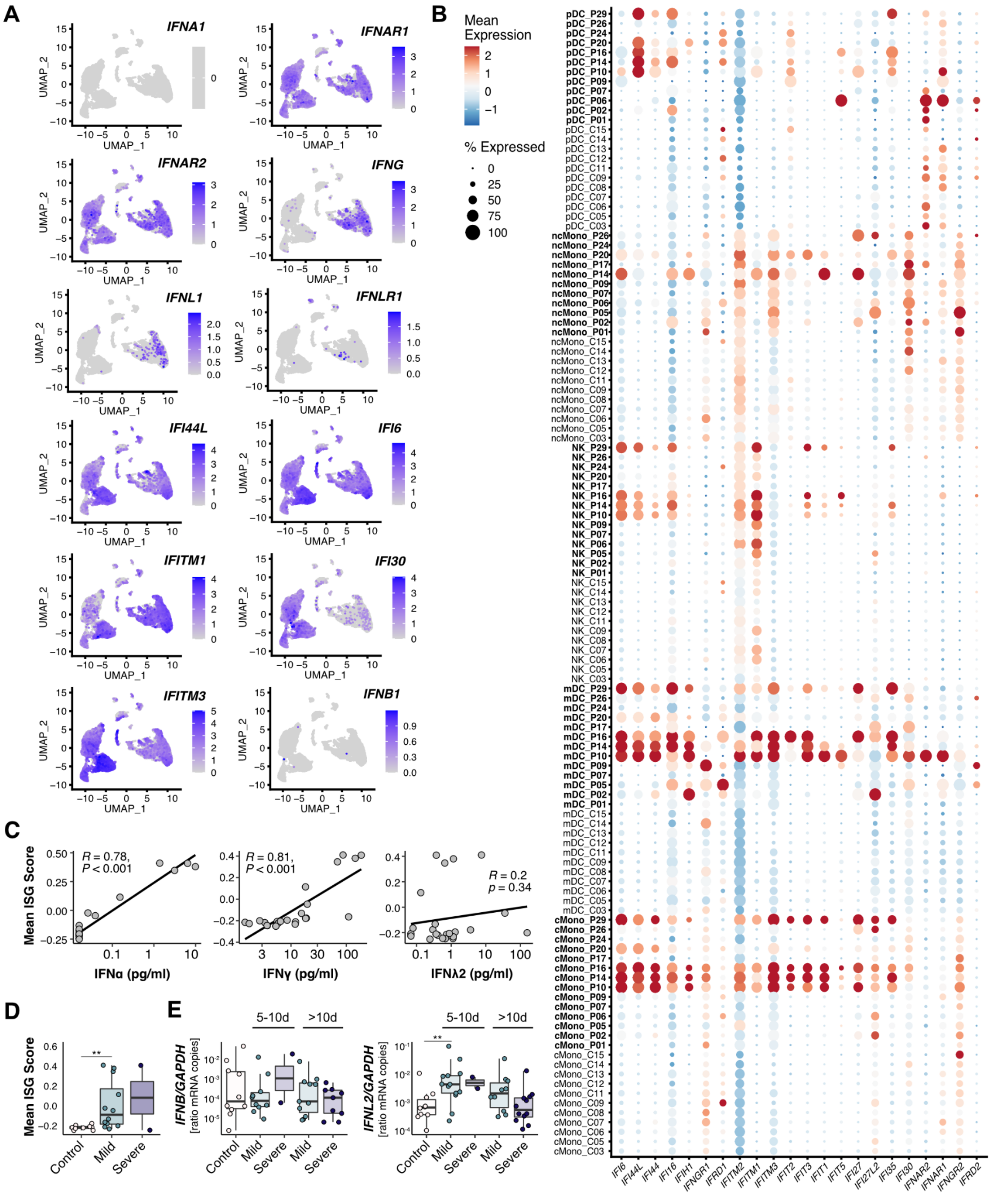


**Figure S3: IFN response in COVID-19 patients, related to Figure 3**

**A)** Expression of IFN-related genes projected onto the UMAP of cell (sub)types of PBMCs identified by scRNAseq from 14 COVID-19 patients and 11 controls. **B)** Dot plots indicating expression of IFN-related genes in different cell types of PBMCs from patients and controls. **C)** Linear regression analysis of the ISG score (as shown in D) versus levels of type I, II, and III IFN levels in plasma. **D)** ISG score (calculated by applying Seurat function *AddModuleScore* to IFN-stimulated genes) in PBMCs from controls and patients with mild and severe COVID-19. **E)** Relative oropharyngeal expression of type I and III IFNs in controls and patients with mild and severe COVID-19 at the early (<10 d after symptom onset) and late (>10 d after symptom onset) phase of infection. See also main text Fig. 3.

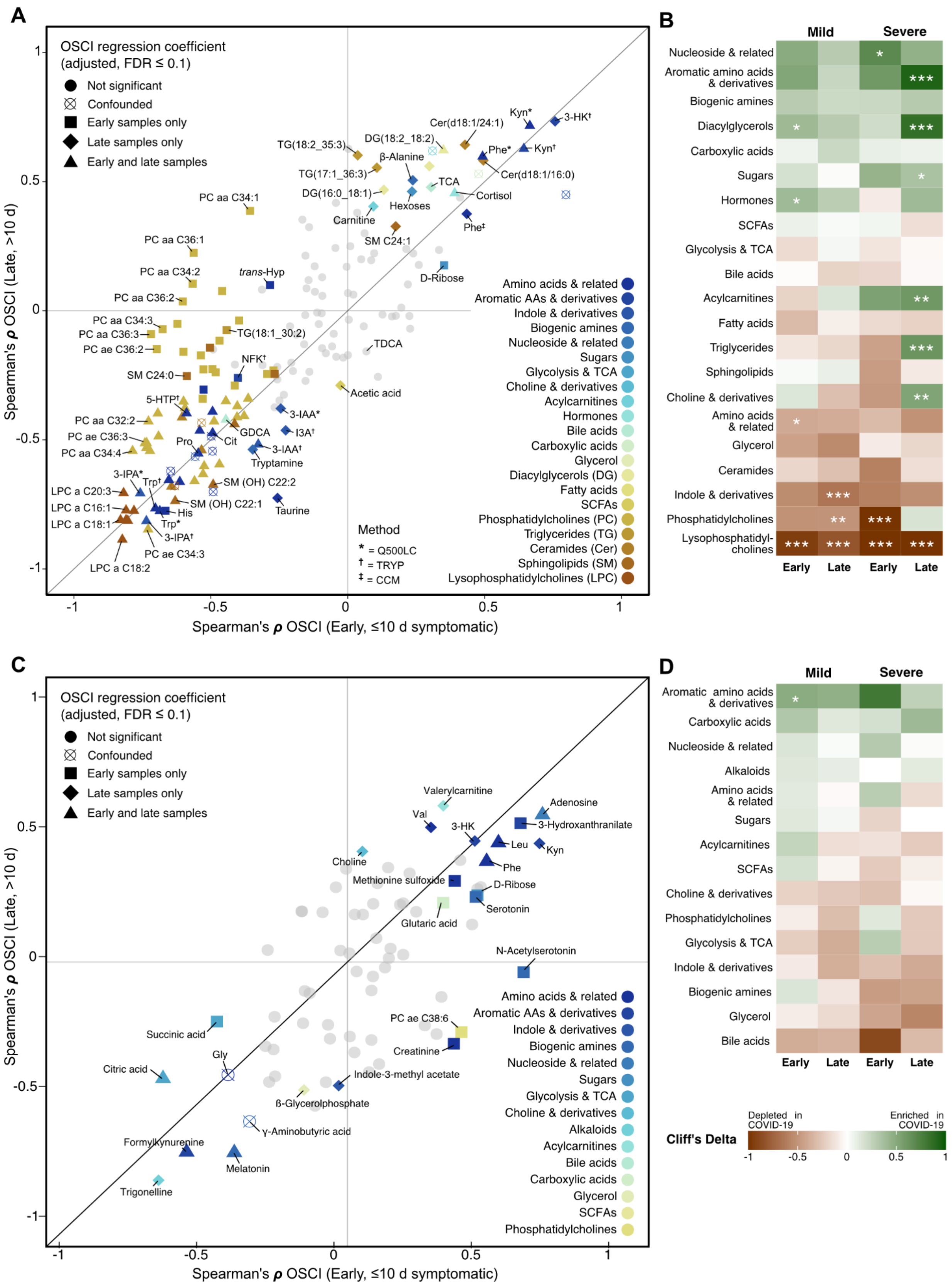
A
OSCI regression coefficient
(adjusted, FDR ≤ 0.1)
Not significant
Confounded
Early samples only
Late samples only
Early and late samples
Kyn*
3-HK†
Cer(d18:1/24:1)
DG(18:2_18:2)
TG(18:2_35:3)
β-Alanine
Phe*
Kyn†
Cer(d18:1/16:0)
TG(17:1_36:3)
TCA
DG(16:0_18:1)
Cortisol
PC aa C34:1
Hexoses
Carnitine
Phe‡
SM C24:1
PC aa C36:1
PC aa C34:2
trans-Hyp
D-Ribose
PC aa C36:2
PC aa C34:3
PC aa C36:3
TG(18:1_30:2)
PC ae C36:2
TDCA
NFK†
SM C24:0
Acetic acid
5-HTP†
3-IAA*
PC aa C32:2
GDCA
I3A†
PC ae C36:3
Pro
Cit
3-IAA†
PC aa C34:4
Tryptamine
3-IPA*
Trp†
SM (OH) C22:2
LPC a C20:3
SM (OH) C22:1
LPC a C16:1
His
Taurine
LPC a C18:1
Trp*
3-IPA†
PC ae C34:3
LPC a C18:2
Amino acids & related
Aromatic AAs & derivatives
Indole & derivatives
Biogenic amines
Nucleoside & related
Sugars
Glycolysis & TCA
Choline & derivatives
Acylcarnitines
Hormones
Bile acids
Carboxylic acids
Glycerol
Diacylglycerols (DG)
Fatty acids
SCFAs
Phosphatidylcholines (PC)
Triglycerides (TG)
Ceramides (Cer)
Sphingolipids (SM)
Lysophosphatidylcholines (LPC)
Method
* = Q500LC
† = TRYP
‡ = CCM
Spearman's ρ OSCI (Late, >10 d)
Spearman's ρ OSCI (Early, ≤10 d symptomatic)
1
0.5
0
-0.5
-1
B
Mild
Severe
Nucleoside & related
Aromatic amino acids & derivatives
Biogenic amines
Diacylglycerols
Carboxylic acids
Sugars
Hormones
SCFAs
Glycolysis & TCA
Bile acids
Acylcarnitines
Fatty acids
Triglycerides
Sphingolipids
Choline & derivatives
Amino acids & related
Glycerol
Ceramides
Indole & derivatives
Phosphatidylcholines
Lysophosphatidylcholines
Early
Late
Early
Late
C
OSCI regression coefficient
(adjusted, FDR ≤ 0.1)
Not significant
Confounded
Early samples only
Late samples only
Early and late samples
Valerylcarnitine
Adenosine
Val
3-HK
3-Hydroxanthranilate
Leu
Kyn
Choline
Phe
Methionine sulfoxide
D-Ribose
Serotonin
Glutaric acid
N-Acetylserotonin
PC ae C38:6
Succinic acid
Creatinine
Gly
Citric acid
Indole-3-methyl acetate
ß-Glycerolphosphate
γ-Aminobutyric acid
Formylkynurenine
Melatonin
Trigonelline
Amino acids & related
Aromatic AAs & derivatives
Indole & derivatives
Biogenic amines
Nucleoside & related
Sugars
Glycolysis & TCA
Choline & derivatives
Alkaloids
Acylcarnitines
Bile acids
Carboxylic acids
Glycerol
SCFAs
Phosphatidylcholines
Spearman's ρ OSCI (Late, >10 d)
Spearman's ρ OSCI (Early, ≤10 d symptomatic)
D
Mild
Severe
Aromatic amino acids & derivatives
Carboxylic acids
Nucleoside & related
Alkaloids
Amino acids & related
Sugars
Acylcarnitines
SCFAs
Choline & derivatives
Phosphatidylcholines
Glycolysis & TCA
Indole & derivatives
Biogenic amines
Glycerol
Bile acids
Early
Late
Early
Late
Depleted in COVID-19
Enriched in COVID-19
Cliff's Delta
-1
-0.5
0
0.5
1

**Figure S4: Metabolite associations with disease severity at both early and late timepoints, related to Figure 4**
**A)** Spearman correlations between plasma metabolite concentrations and OSCI scores at the time of sampling were calculated using either early (N=26) or late (N=40) subsets of COVID-19 patient samples and healthy controls (N=30), and are shown contrasted with one another. Metabolites are annotated to show their biochemical class and analytical method. Shapes were assigned according to the integrated output from our modeling with clinical variables in each subset (early and late) separately, after adjusting for case-control disease status. Compounds closer to the diagonal line displayed consistent severity correlations regardless of sample timepoint, while levels of several phosphatidylcholines were negatively correlated with disease severity in early samples only. **B)** Cliff's delta effect sizes for different subsets of COVID-19 patients compared to controls for each chemical class. Adjusted significances from Mann-Whitney U tests which were not confounded from our modeling analysis with clinical covariates are depicted (FDR<0.1=*, FDR<0.01=**, FDR<0.001=***). **C-D)** The same as in A) and B) except for urine metabolites. See also main text Fig. 4. Phe: phenylalanine, Pro: proline, Cit: citrullin, *trans*-Hyp: *trans*-4-hydroxyproline, Trp: tryptophan, Kyn: kynurenine, 5-HTP: 5-hydroxytryptophan, 3-HK: 3-hydroxykynurenine, NFK: N-Formylkynurenine, TDCA: taurodeoxycholic acid, GDCA: glycodeoxycholic acid, TCA: taurocholic acid, 3-IPA: indole-3-propionic acid, 3-IAA: indole-3-acetic acid, I3A: indole-3-carboxaldehyde; HAP: hospital-acquired pneumonia

The following tables will be uploaded separately for publication but are currently available at www.github.com/sxmorgan/pa-covid-multi-omics.

**Table S1: Cohort characteristics, related to Table 1**

**Table S2: Clinical lab parameters, related to Figure 1**

**Table S3: Taxonomic profiles, related to Figures 2, S2**

**Table S4: Immune response profiles, related to Figures 3, S3**

**Table S5: Metabolite profiles, related to Figures 4, S4**

**Table S6: Results from statistical analysis with clinical variables, related to Figures 2-4, S2-S4**

**Table S7: Integrated statistical analysis results, related to Figure 5**

**Table S8: Quality Control reporting of metabolomics data according to mQACC standards**

**Table S9: Ultra-high performance liquid chromatography (UHPLC) gradient for LC part**

| | LC1 | | | | LC2 | | | |
|---|---|---|---|---|---|---|---|---|
| No | Time [min] | Flow [mL/min] | A [%] | B [%] | Time [min] | Flow [mL/min] | A [%] | B [%] |
| 1 | 0.00 | 0.8 | 100 | 0 | 0.00 | 0.8 | 100 | 0 |
| 2 | 0.25 | 0.8 | 100 | 0 | 0.25 | 0.8 | 100 | 0 |
| 3 | 1.50 | 0.8 | 88 | 12 | 0.50 | 0.8 | 75 | 25 |
| 4 | 2.70 | 0.8 | 82.5 | 17.5 | 2.00 | 0.8 | 50 | 50 |
| 5 | 4.00 | 0.8 | 50 | 50 | 3.00 | 0.8 | 25 | 75 |
| 6 | 4.50 | 0.8 | 0 | 100 | 3.50 | 0.8 | 0 | 100 |
| 7 | 4.70 | 1.0 | 0 | 100 | 4.70 | 1.0 | 0 | 100 |
| 8 | 5.00 | 1.0 | 0 | 100 | 5.00 | 1.0 | 0 | 100 |
| 9 | 5.10 | 1.0 | 100 | 0 | 5.10 | 1.0 | 100 | 0 |
| 10 | 5.80 | 1.0 | 100 | 0 | 5.80 | 1.0 | 100 | 0 |

**Table S10: Ultra-high performance liquid chromatography (UHPLC) gradient for flow injection (FIA) part**

| No | Time [min] | Flow [mL/min] | A [%] | B [%] |
|---|---|---|---|---|
| 1 | 0.0 | 0.03 | 0 | 100 |
| 2 | 1.6 | 0.03 | 0 | 100 |
| 3 | 2.4 | 0.20 | 0 | 100 |
| 4 | 2.8 | 0.20 | 0 | 100 |
| 5 | 3.0 | 0.03 | 0 | 100 |

**Table S11: Acquisition method parameters for liquid chromatography (LC) and flow injection analysis (FIA)**

| Option | Parameter | LC1 | LC2 | FIA1 | FIA2 |
|---|---|---|---|---|---|
| MS | Scan type | MRM | MRM | MRM | MRM |
| | Polarity | Positive | Negative | Positive | Negative |
| | MRM detection window (sec) | 30 | 30 | - | - |
| | Duration (min) | 5.45 | 5.45 | 2.95 | 2.95 |
| | Delay time (sec) | 0 | 0 | 0 | 0 |
| | Cycle (sec) | 0.25 | 0.15 | N/A | N/A |
| Advanced MS | Resolution Q1 | Unit | Unit | Unit | Unit |
| | Resolution Q3 | Unit | Unit | Unit | Unit |
| | Intensity threshold | 0 | 0 | 0 | 0 |
| | Setting time (ms) | 0 | 0 | 0 | 0 |
| | Pause between mass ranges (ms) | 2 | 2 | 5.007 | 3 |
| Source/ gas | Curtain gas | 45 | 20 | 20 | 10 |
| | Collision gas | 9 | 8 | 9 | 9 |
| | Ion spray voltage | 5500 | -4500 | 5500 | 5500 |
| | Temperature | 500 | 650 | 200 | 350 |
| | Ion source gas 1 | 60 | 40 | 40 | 30 |
| | Ion source gas 2 | 70 | 40 | 50 | 85 |

MS: mass spectrometry

**Table S12: List of metabolite derivatives and their biological groups used for reference search**

| Group | Metabolite | Abbreviation | Detected as |
|---|---|---|---|
| AA | Alanine | Ala | 3TMS |
| | | | 2TMS |
| AA | Asparagine | Asn | 2TMS |
| AA | Aspartic acid | Asp | 2TMS |
| | | | 3TMS |
| AA | Cysteine | Cys | 3TMS |
| AA | Glycine | Gly | 2TMS |
| | | | 3TMS |
| AA | Isoleucine | Ile | 1TMS |
| | | | 2TMS |
| AA | Leucine | Leu | 1TMS |
| | | | 2TMS |
| AA | Lysine | Lys | 3TMS |
| AA | Methionine | Met | 1TMS |
| | | | 2TMS |
| AA | Phenylalanine | Phe | 1TMS |
| | | | 2TMS |
| AA | Proline | Pro | 1TMS |
| | | | 2TMS |
| AA | Serine | Ser | 2TMS |
| | | | 3TMS |
| | | | 4TMS |
| AA | Threonine | Thr | 2TMS |
| | | | 3TMS |
| AA | Tryptophan | Trp | 2TMS |
| AA | Tyrosine | Tyr | 3TMS |
| AA | Valine | Val | 1TMS |
| | | | 2TMS |
| Glycerol | Dihydroxyacetone phosphate | DHAP | 1MeOX 3TMS |
| Glycerol | Glycerol | Glyc | 3TMS |
| Glycerol | Glycerol-3-phosphate | Glyc3P | 4TMS |
| Glycolysis | Fructose-6-phosphate | F6P | 1MeOX 6TMS |

| | | | |
|---|---|---|---|
| Glycolysis | Glucose-6-phosphate | G6P | 1MeOX 6TMS |
| Glycolysis | Glyceric acid-3-phosphate | GA3P | 4TMS |
| Glycolysis | Lactic acid | Lac | 2TMS |
| Glycolysis | Phosphoenolpyruvic acid | PEP | 3TMS |
| Glycolysis | Pyruvic acid | Pyr | 1MeOX 1TMS |
| Nucleobase | Adenine | Adenine | 2TMS |
| Nucleobase | Uracil | Uracil | 2TMS |
| Nucleoside | Adenosine | Adenosine | 3TMS |
| | | | 4TMS |
| Nucleoside | Cytosine | Cytosine | 2TMS |
| Others | Butanoic acid, 3-hydroxy- | But3h | 2TMS |
| Others | Butanoic acid, 4-amino- | But4am | 3TMS |
| Others | Erythritol | Ery | 4TMS |
| Others | Glutaric acid | Glut | 2TMS |
| Others | Glyceric acid | Glyc | 3TMS |
| PPP | Ribose-5-phosphate | R5P | 1MeOX 5TMS |
| PPP | Ribose | Ribose | 1MeOX 4TMS |
| TCA | Citric acid | Cit | 4TMS |
| TCA | Fumaric acid | Fum | 2TMS |
| TCA | Glutaric acid, 2-hydroxy- | 2HG | 3TMS |
| TCA | Glutaric acid, 2-oxo- | aKG | 1MeOX 2TMS |
| TCA | Malic acid | Mal | 3TMS |
| TCA | Succinic acid | Suc | 2TMS |

AA: Amino acids. PPP: Pentose phosphate pathway. TCA: Tricarboxylic acid cycle. TMS: Trimethylsilyl derivatives. MeOX: Methoxyamine hydrochloride
